# An amphipathic helix facilitates direct membrane binding of Mycoplasma FtsZ

**DOI:** 10.1101/2023.08.29.555414

**Authors:** Soumyajit Dutta, Sakshi Poddar, Joyeeta Chakraborty, Ramanujam Srinivasan, Pananghat Gayathri

## Abstract

Cell division in bacteria is initiated by constriction of the Z-ring comprising two essential proteins FtsZ and FtsA. Despite our knowledge about the crucial function of the Z-ring in bacterial division, the precise roles and mechanism of how FtsZ and FtsA drive cell constriction remain elusive. FtsZ/FtsA in wall-less bacteria like mycoplasmas is an ideal model system for obtaining mechanistic insights into Z-ring constriction in the absence of cell wall machinery. In this study, we have analyzed FtsZ and FtsA sequences of 113 mycoplasma species and compared with the corresponding protein sequences in cell-walled bacteria. We report a phylogenetically distinct group of 12 species that possess FtsZs without the canonical FtsA interacting conserved C-terminal peptide (CCTP) motif. Interestingly, these FtsZs contain a putative membrane-binding amphipathic helix as an N-terminal or C-terminal extension to the globular FtsZ domain. As a proof-of-concept, we experimentally show that the proposed C-terminal amphipathic helix in *M. genitalium* FtsZ binds liposomes *in vitro* as well as localizes to *E. coli* membrane *in vivo*. Additionally, we identify a putative cholesterol recognition motif within the C-terminal amphipathic helix region of *M. genitalium* FtsZ. Our study catalogues the functional variations of membrane attachment by the FtsZ and FtsA system in cell wall-less mycoplasmas and provides a new perspective to study novel functions of FtsZ/A system in cell division.

**Importance:** Z-ring and peptidoglycan synthesis machinery both play crucial roles in bacterial cell division. Currently, our knowledge about how FtsZ and FtsA, the two primary components of the Z-ring, function, is limited to cell-walled bacteria where ring constriction is coupled to peptidoglycan synthesis. Cell wall-less bacterial FtsZ/A system is an excellent model to study the mechanism of Z-ring constriction in the absence of cell wall synthesis machinery. Here, we analysed FtsZ protein sequences across mycoplasma species and identified their characteristic sequence features. Our study reveals a novel group of FtsZs from mycoplasma with an inherent membrane binding and probable cholesterol sensing amphipathic motif, which serves as a new paradigm to explore fundamental roles of FtsZ and FtsA in Z-ring constriction during bacterial division.

## Introduction

FtsZ and FtsA are the two major proteins involved in bacterial cell division (1, 2). The tubulin-like protein FtsZ forms short patches of overlapping filaments called the Z-ring which treadmills and gradually leads to constriction of the cell (3, 4). The presence of FtsZ is essential to initiate constriction but seems to be dispensable at the later stages of bacterial cell division (5, 6). Treadmilling FtsZ filaments act as a guide for incorporating peptidoglycan strands in the newly forming septum during constriction (3, 4, 7, 8). In most bacteria, the actin-homolog FtsA, with the help of its C-terminal amphipathic helix, functions as the primary membrane anchor for FtsZ (9, 10). FtsZ and FtsA interact with other cell division-related proteins to further regulate the downstream events in the constriction process (11–15). Other than FtsA, there are proteins such as ZipA (in Gram-negative gamma proteobacteria) and SepF (in Gram-positive bacteria and cyanobacteria) that act as membrane anchors for FtsZ (16, 17).

The domain organization of FtsZ, a tubulin-like protein, contains a GTP binding globular core, an N-terminal extension, a C-terminal disordered linker, and a conserved C-terminal peptide (CCTP) (Fig.1b) (18). The globular core domain contains two highly conserved tubulin signature motifs, (1) GTP binding motif-[S/A/G]–G–G–T–G–[S/A/T]–G, and (2) GTP hydrolysis motif-N–X–D–X–X–D, (where X can be any residue) (19, 20). The CCTP motif also shows a high degree of conservation across different bacterial species with a consensus sequence of D/E–I/V–P–X–F/Y–L–R/K, (where X can be any residue) (21, 22). The CCTP is required for interaction between FtsZ and many of its interacting membrane tethering proteins like FtsA, and the less conserved ZipA or SepF (23–25).

FtsA acts as the primary membrane anchor for FtsZ and it is widely conserved in most bacterial species. FtsA is an actin family protein containing a globular core domain followed by a disordered linker and C-terminal amphipathic helix (CTAH) (Fig. 1b). Membrane binding ability of the CTAH of FtsA helps FtsZ filaments assemble at the inner cytoplasmic membrane and initiate divisome formation (9, 26). However, the precise role of this essential protein in the divisome assembly is not clear yet. Although protofilament architecture of FtsA has been captured *in vitro* for *Thermotoga maritima* (27), *Streptococcus pneumoniae* (28, 29), *Escherichia coli* (30, 31), and *Vibrio cholerae* (32), the ATPase activity of *T. maritima* or *S. pneumoniae* has not been observed despite the presence of ATPase domain. Rapid ATP hydrolysis for *E. coli* FtsA has been observed in presence of phospholipids (33) while *in vitro* Z-ring dynamics does not require ATPase activity of FtsA (34). Thus, while a passive membrane anchoring role is important, it is not yet clear if FtsA plays additional active role through ATP hydrolysis.

**Fig. 1.**
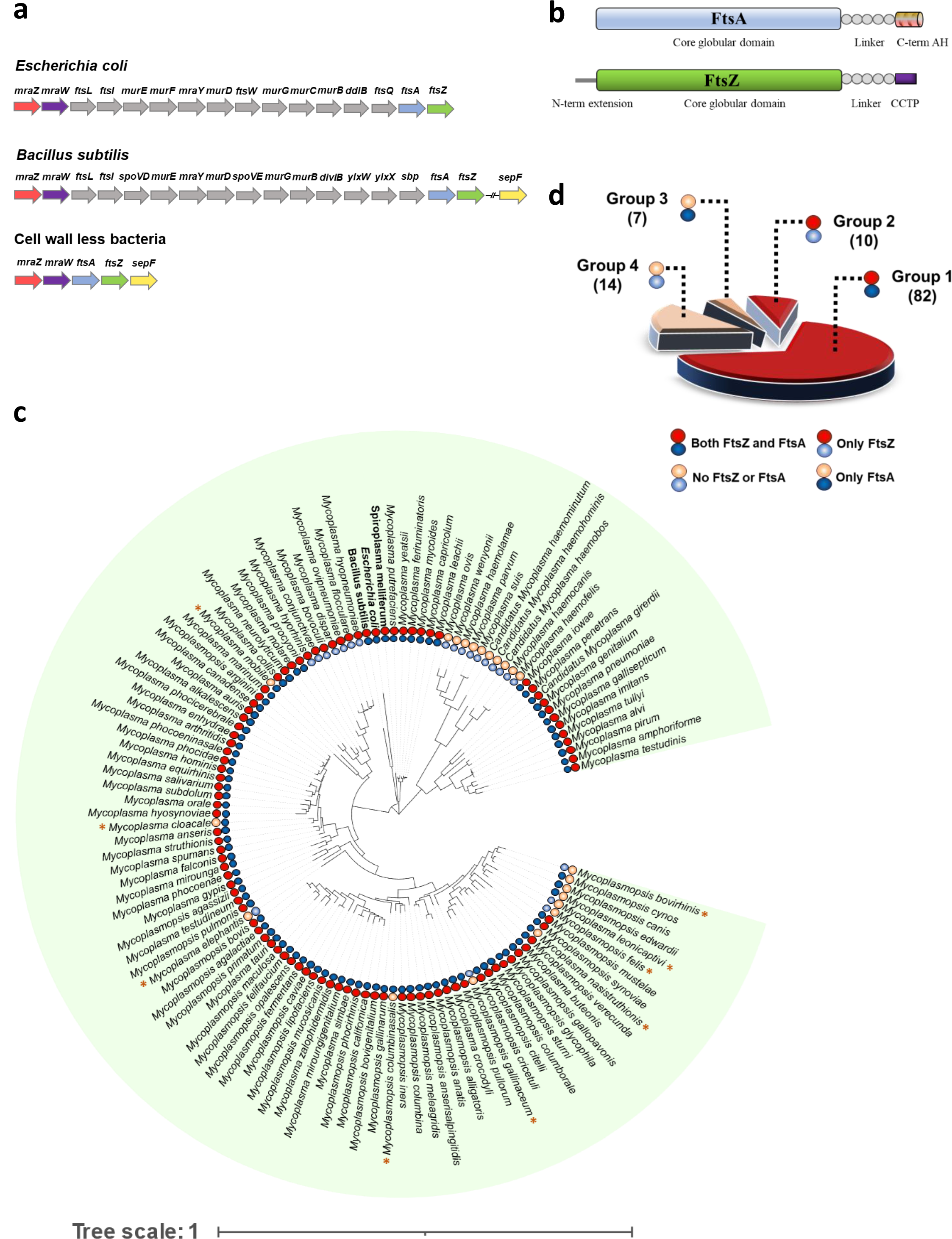
A scheme for classification of mycoplasma species based on the presence of *ftsA* and *ftsZ* in their genomes. (a) Genomic locus of division and cell wall cluster (*dcw*) operon genes of *Escherichia coli*, *Bacillus subtilis,* and cell wall-less bacteria. The arrowhead indicates gene orientation 5′ → 3′. (b) Domain architecture of FtsA, FtsZ. (c) 16S rRNA-based tree of 113 mycoplasma species is constructed using maximum likelihood analysis and the distribution of FtsZ and FtsA proteins are highlighted. Solid red (for FtsZ) and solid blue (for FtsA) circles represent the presence of the proteins and transparent colored circles indicate their absence. Outlier species which are away from clusters of the respective groups are marked as yellow colored star (*). (d) Grouping of mycoplasmas based on the presence of FtsZ and FtsA. The number of organisms consisting each group is mentioned within brackets.

Cell division is a complex process that relies on coordinated and collective action of a number of proteins involved in Z-ring assembly, cell wall synthesis and their regulations (35). Overall, in bacteria, there are at least twenty different proteins that play an important role in the formation of divisome, a multiprotein complex that coordinates different steps of the division in a highly regulated and organized manner (36). Notably, very often in bacteria, many of the division-related proteins are transcribed from a single operon called division and cell wall (*dcw*) cluster (37). The *dcw* operon in cell walled model organisms *E. coli* and *Bacillus subtilis* comprises sixteen and seventeen genes, respectively, which include FtsZ, FtsA and their downstream interactor proteins involved in cell wall synthesis (Fig. 1a). While much of the information regarding the function of cell division proteins has been deciphered based on studies carried out in walled bacterial species, our understanding of cell wall-less systems is limited. In cell wall-less bacteria, genes related to peptidoglycan synthesis are lost and a concomitant shorter *dcw* operon is the result of an extensive genome reduction (38). In mycoplasmas, for example, only five genes namely *mraZ*, *mraW (rsmH)*, *ftsA, ftsZ* and *sepF* can be found in the *dcw* operon (39, 40). The highly conserved nature of these genes across the bacterial kingdom implies their important function in cell division in a cell wall less milieu. In cell-walled bacteria, cell wall synthesis by the peptidoglycan synthesis machinery is supposedly the major player for constriction force generation as the inward force generated by the Z-ring is not sufficient to overcome the turgor pressure (7). However, how constriction occurs in cell wall-less bacteria like mycoplasmas, where FtsZ and FtsA are possibly the only two proteins to constitute the Z-ring in the absence of the cell wall synthesis machinery, is an enigma.

Earlier studies on *Mycoplasma genitalium* suggest that the cell is capable of division in the absence of FtsZ without affecting its doubling rate (41). In *M. pneumoniae*, a close relative of *M. genitalium*, motility-driven mechanical force appears to facilitate constriction, a mechanism alternatively termed as cytofission (42). Other cell wall-less strains of bacteria including the L-form have been found to utilize processes such as membrane blebbing, budding, or extrusion-resolution for multiplying (43, 44). Thus, if FtsZ is not essential, why it remains conserved in genome of cell wall-less bacteria, remains unexplained. However, in *M. genitalium,* FtsZ has been found to localize at the mid cell at the onset of constriction indicating its role in cell division (45). Interestingly, a recent discovery on mycoplasma JCVI-syn3a, the minimal cell with synthetic genome, shows the requirement *ftsZ* and *sepF* for normal cell morphology and division (40). This essentially makes all five *dcw* cluster genes namely *mraZ*, *mraW*, *ftsA*, *ftsZ*, and *sepF* an integral part of the minimal gene set in JCVI-syn3a. These findings prompted us to look into characteristic sequence features of these cell division proteins across mycoplasma species to understand their role in cell wall-less bacteria.

Here we report the sequence features of the primary contractile ring forming protein FtsZ across 113 mycoplasma species. We identified a phylogenetically distinct FtsZ cluster comprising 12 mycoplasmas having an amphipathic helix with potential membrane binding characteristics. *M. genitalium*, the bacterium with a minimal genome, is one of the member species of this clade, which interestingly, possesses a C-terminal amphipathic helix (CTAH). CTAH will possibly allow a direct membrane attachment for FtsZ bypassing the requirement of FtsA as membrane anchor. Furthermore, we have experimentally shown that the *M. genitalium* CTAH motif binds to liposomes *in vitro* as well as localizes to *E. coli* membrane *in vivo*. We also identified a cholesterol recognition amino acid consensus (CRAC) sequence in the C-terminal of *M. genitalium* and *M. pneumoniae* FtsZ. Overall, our identification of unique membrane binding and cholesterol interacting properties of bacterial FtsZ provide us with examples of naturally existing minimal divisome components, a powerful handle to obtain new insights to the fundamental mechanism of cell division.

## Results

### All mycoplasma species do not possess *ftsZ* and *ftsA* genes

We analysed 113 mycoplasma genomes in NCBI database for division and cell wall (*dcw*) operon to check the presence of division related genes (Supplementary data sheet). While searching the NCBI genome database, we found *ftsA* as an unannotated gene present in between *mraW* and *ftsZ* in most mycoplasmas. After checking the domain organization of its protein coding sequence, we confirmed this as FtsA (Fig. S1). Importantly, in mycoplasma, the four division related genes namely *mraZ*, *mraW*, *ftsA*, and *ftsZ* were present in the same sequential manner as found in cell walled bacteria. As expected, all additional genes related to the cell wall synthesis present in the *dcw* operon in cell-walled bacteria were absent. We observed that even FtsZ and FtsA were not conserved across all mycoplasma species. Accordingly, based on the presence of FtsA and FtsZ, we divided them into four groups (Fig. 1c, d). Based on the comparison with the 16S rRNA based phylogenetic tree, we observed that the presence of FtsZ/FtsA was correlated to the species organization, except for a few species that were outliers (highlighted by ‘*’; Fig. 1c, d).

Majority of them i.e., 82 out of 113 species, were classified into group 1 where both FtsZ and FtsA are present (Fig. 1d). Group 2, having only FtsZ but no FtsA, includes 10 species while group 3 has 7 species where FtsA is present but FtsZ is absent. Finally, the 14 species of group 4 lack both FtsZ and FtsA. Further, we analysed the sequences systematically to find out the characteristic features of these two proteins in the various groups and to further correlate with their role in cell division.

### FtsZ sequence analysis reveals variable C-terminal sequences among mycoplasma species

To compare the FtsZ proteins among group 1 mycoplasma species, we checked for the conservation of three characteristic signature motifs of FtsZ: the GTP binding motif, GTPase motif and CCTP (18, 27). We found the GTP binding and hydrolysis motifs were well conserved across species whereas the CCTP was variable (Fig. 2a). We divided the sequences into five subgroups based on the signature amino acid sequences of FtsZ CCTP (Fig 2a, S2). The cladogram constructed from the FtsZ sequences interestingly matched with the CCTP based classification (Fig. 2b). The CCTP based groups of FtsZ also forms clusters in the 16S rRNA based cladogram (Fig. S3. 14 organisms out of 82 possessed a canonical FtsZ CCTP and were classified into group 1A. Sequences that showed conservation of charges similar to canonical CCTP residues, but missed the conserved proline, were classified into group 1B. Group 1C sequences were more divergent where only proline and phenylalanine were conserved as found in cell walled bacterial CCTPs while the negatively charged stretch of residues were not conserved. The rest of the species did not show any conservation in their CCTP and while comparing with the cladogram we identified two such clusters (called 1D and 1E) separated by group 1C. Multiple sequence alignment of the C-terminal FtsZs of 1D and 1E groups showed conservation of negative charge and some hydrophobic residues (Fig. S2), though not belonging to the canonical motif signature. Importantly, the invariant proline of CCTP is missing in groups 1B, 1D and 1E. We also report an extended N-terminal region (i.e. more than 15 amino residues compared to the core globular domain of cell walled bacteria FtsZs) in groups D and E (except *M. genitalium* and *M. pneumoniae*) FtsZ sequences (Fig. S4, Table S1).

**Fig. 2.**
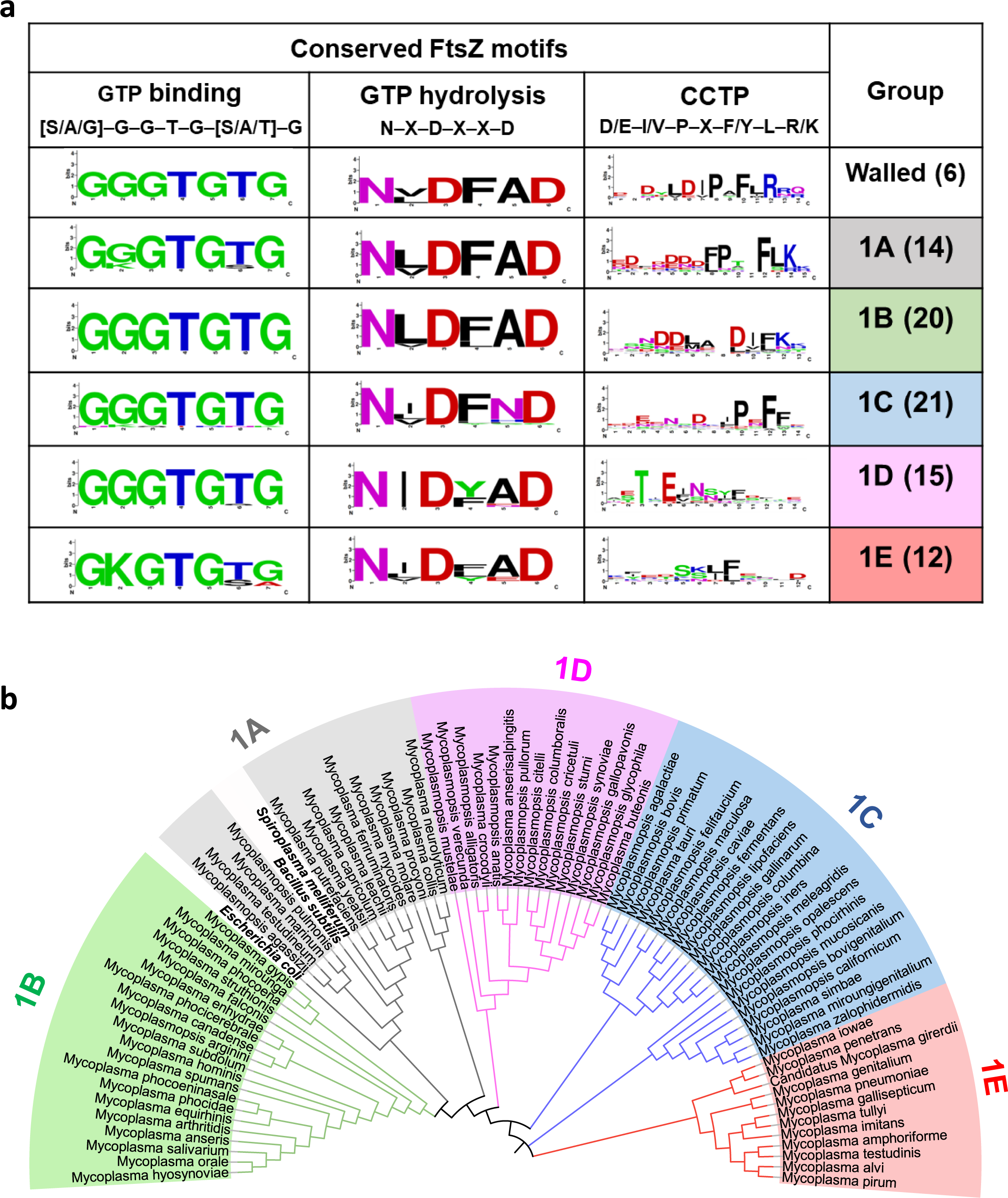
Classification based on conserved C-terminal peptide (CCTP) sequences matches with the phylogenetic relationship of mycoplasma FtsZs. (a) Group 1 mycoplasmas having FtsZ and FtsA are grouped into five categories based on the various types of CCTP present. Groupwise WebLogos are shown for three FtsZ motifs: GTP binding, hydrolysis, and CCTP which are found to be conserved in cell-walled bacteria. The number within the bracket for a group represents number of sequences used for generating the Weblogo. (b) Cladogram of FtsZ from group 1A mycoplasma species is constructed using the maximum likelihood method. Different clusters are highlighted and labeled as 1A – E as per our annotated groups based on CCTP motifs presents at the C-terminal end of FtsZ.

### Mycoplasma FtsZs of Group 1E possess an amphipathic helix

One of the interesting findings of our analysis is the presence of a potential amphipathic helix (AH) in each of the twelve FtsZ sequences of group 1E, where CCTP motifs for FtsA interaction are probably absent. Ten FtsZ sequences were identified to possess AH at the N-terminal and the other two (*M. genitalium* and *M. pneumoniae*) at the C terminal (Fig. S5). The helical wheel diagram highlights the amphipathic nature of the proposed N-terminal and C-terminal helical regions (Fig. 3a, Table S2). We also calculated two parameters related to amphipathic helix, namely, hydrophobicity and hydrophobic moments for all the proposed AHs and compared with known membrane binding AHs from FtsA, SepF and MinD from cell walled bacteria. Importantly, the amphipathic parameters for all the twelve proposed mycoplasma AHs fall in the same region indicating their probable amphipathic membrane binding nature (Fig. 3b).

**Fig. 3.**
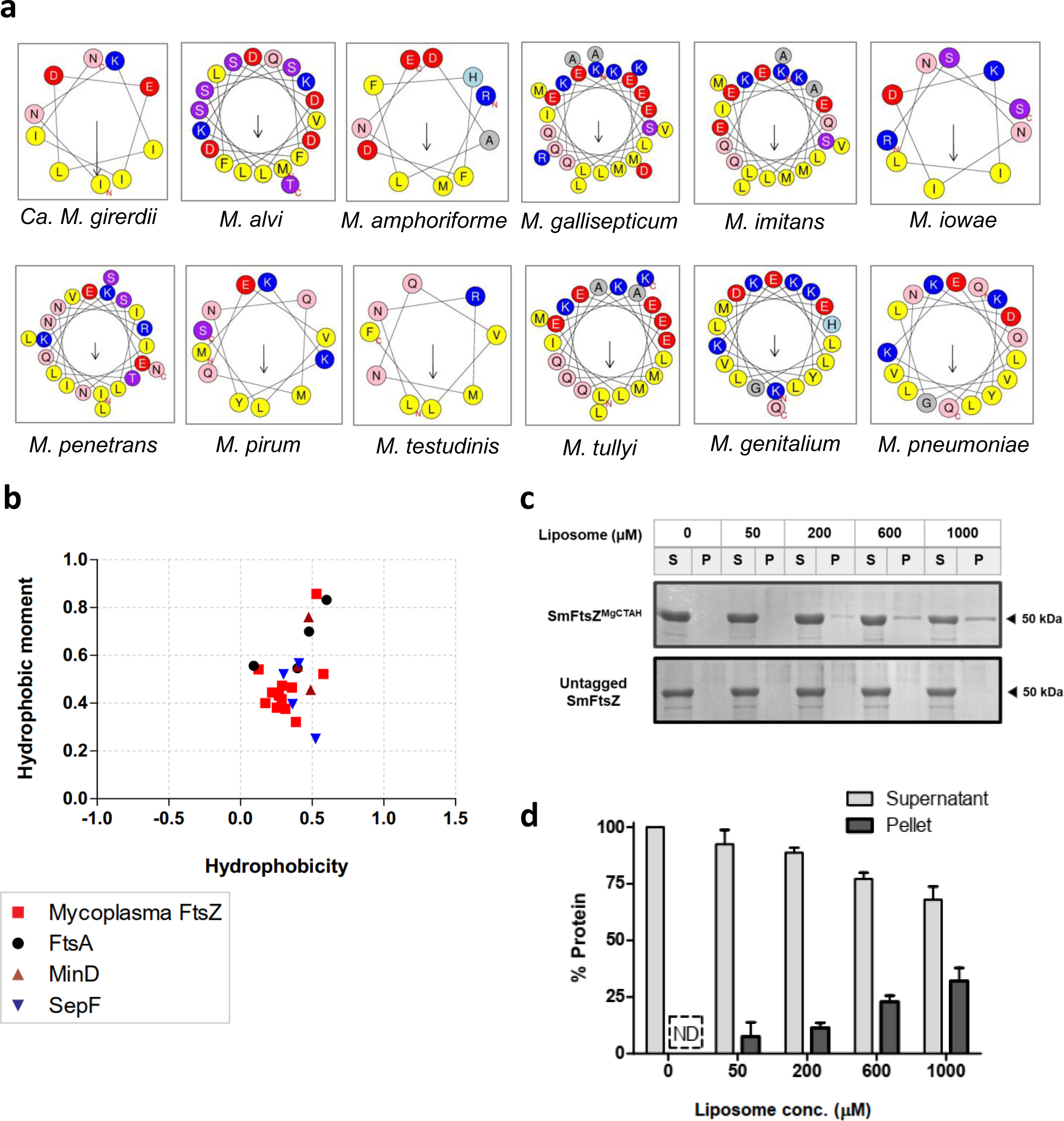
Mycoplasma FtsZ representative of Group 1E binds to membrane directly through an inherent amphipathic helix. (a) Helical wheel projections show the amphipathic nature of the helix-forming amino acids at the FtsZ terminals. Hydrophobic residues are depicted in yellow showing the amphipathic face. Arrows indicate the hydrophobic moments of the respective helices. (b) Hydrophobic moment plot of the putative N-terminal and C-terminal α-helices of group 1E FtsZs. The hydrophobic moment is plotted against hydrophobicity to predict its potential to bind to the membrane. Known amphipathic helices from FtsA, MinD, and SepF also plotted simultaneously for comparison. (c) A representative 12% SDS-PAGE gel showing the presence of SmFtsZ^MgCTAH^ and Untagged SmFtsZ protein in the supernatant (S) and pellet fraction (P) with increasing concentrations of liposomes from 0 µM (i.e. no liposome control) to 1000 µM. (d) Quantification plot for SmFtsZ^MgCTAH^ protein in the pellet fraction. The fraction of protein was calculated from the SDS PAGE gel band intensity in the pellet fraction divided by the sum of band intensities in pellet and supernatant. Error bar denotes standard deviation calculated from three independent experiments. ND: Not detectable.

### Membrane binding features of Mycoplasma FtsAs and SepFs show varied combinations of membrane attachment

The lack of canonical CCTP motif and the presence of an inherent amphipathic helix in FtsZs prompted us to check whether an amphipathic helix is present in the corresponding group 1E FtsAs. The amphipathic nature of the FtsA C-terminal helix was checked by plotting hydrophobic moment and hydrophobicity and compared with known membrane binding amphipathic helix. We found that all FtsA sequences from 1E group (except *M. iowae*) possess the canonical C-terminal amphipathic helix (Fig. S6a, b). Moreover, when we checked FtsA sequences from other groups, we found organisms from group 1A and group 1B without the C-terminal amphipathic helix which interestingly form separate clusters in the phylogenetic tree (Fig. S6a). We next checked for the presence of the alternative membrane attachment adaptor SepF and found that it is present in the genome of only six mycoplasma species of group 1A and its locus is next to the *ftsZ* gene of the *dcw* cluster (Fig. S6c). Interestingly, five out of six SepF containing species lack the C-terminal amphipathic helix in the FtsA (Fig. S6d). We also checked for potential membrane binding site at the N-terminal end of the six mycoplasma SepF sequences by helical wheel diagram. Though, no specific hydrophobic face or amphipathic helix was identified similar to that of *B. subtilis* SepF (Fig. S7 a, b), there are conserved multiple hydrophobic amino acids (e.g. phenylalanine, tryptophan etc.) which might be responsible here for hydrophobic loop mediated membrane binding (Fig. S7b). Thus, FtsZs in mycoplasma groups can anchor to the membrane in a variety of modes starting with its inherent amphipathic helix, or through FtsA amphipathic helix/hydrophobic loop as well as through SepF where FtsA AH is absent.

### *M. genitalium* FtsZ CTAH binds liposomes and localises to membranes in *E. coli*

To experimentally verify the membrane binding ability of the proposed AHs, we chose *M. genitalium* sequence as a representative FtsZ with amphipathic helix at the C-terminal end. The proposed amphipathic helix from *M. genitalium* FtsZ i.e. the C-terminal 23 amino acid stretch (347-366) was inserted at the C-terminal end of *Spiroplasma melliferum* (Sm) FtsZ protein to obtain a chimeric construct (named as SmFtsZ^MgCTAH^). We used liposomes made with polar lipids (in molar ratio of 25% DOPG, 20% DOPC, 30% cardiolipin, and 25% spingomyelin) according to their relative abundance in mycoplasma membrane (46). Liposome sedimentation assay confirmed that SmFtsZ^MgCTAH^ binds liposomes as we could see the presence of protein in the pellet fraction which increases with liposome concentration (Fig. 3c, d). In contrast, the untagged SmFtsZ protein (where instead of *M. genitalium* FtsZ CTAH motif, *S. melliferum* FtsZ CCTP motif is present) did not bind to liposomes as no detectable protein band was observed in the pellet of liposomes (Fig. 3c).

Further, we tested if the predicted C-terminal amphipathic helix could localise the core region (amino acids 1 – 366 lacking the CCTP) of EcFtsZ to membranes in *E. coli*. Earlier studies have shown that the core EcFtsZ_1-366_-mVenus fusions localise to the membranes and organise into spiral polymers or ring-like structures only in the presence of a membrane anchor such as the amphipathic helix (residues 254 – 270) from EcMinD. We thus created a translation fusion of the core region of EcFtsZ_1-366_-mNeonGreen (EcFtsZ_1-366_-mNG) to the MinD amphipathic helix (EcFtsZ_1-366_-mNG-MinD^AH^) or the CTAH from *M. genitalium* FtsZ (EcFtsZ_1-366_-mNG-MgFtsZ^CTAH^) and tested the localisation in an *E. coli* strain (JKD7-1) depleted of the native endogenous FtsZ. At 30 °C (in the presence of the native full-length endogenous FtsZ), cells showed normal morphology and both EcFtsZ_1-366_-mNG-MinD^AH^ and EcFtsZ_1-366_-mNG-MgFtsZ^CTAH^ localised to Z-rings, possibly due to co-polymerization with the endogenous full-length FtsZ (Fig. 4a, b). However, at 42°C, when the cells appear filamentous (due to depletion of endogenous FtsZ), EcFtsZ_1-366_-mNG-MgFtsZ^CTAH^ was seen localised to the membranes (Fig. 4d) and was similar to those formed by EcFtsZ_1-366_-mNG-MinD^AH^ (Fig. 4c). We thus conclude that the predicted 23 amino acid residues at the C-terminus of MgFtsZ can function as an amphipathic helix and target localization to liposomes and membranes.

**Fig. 4.**
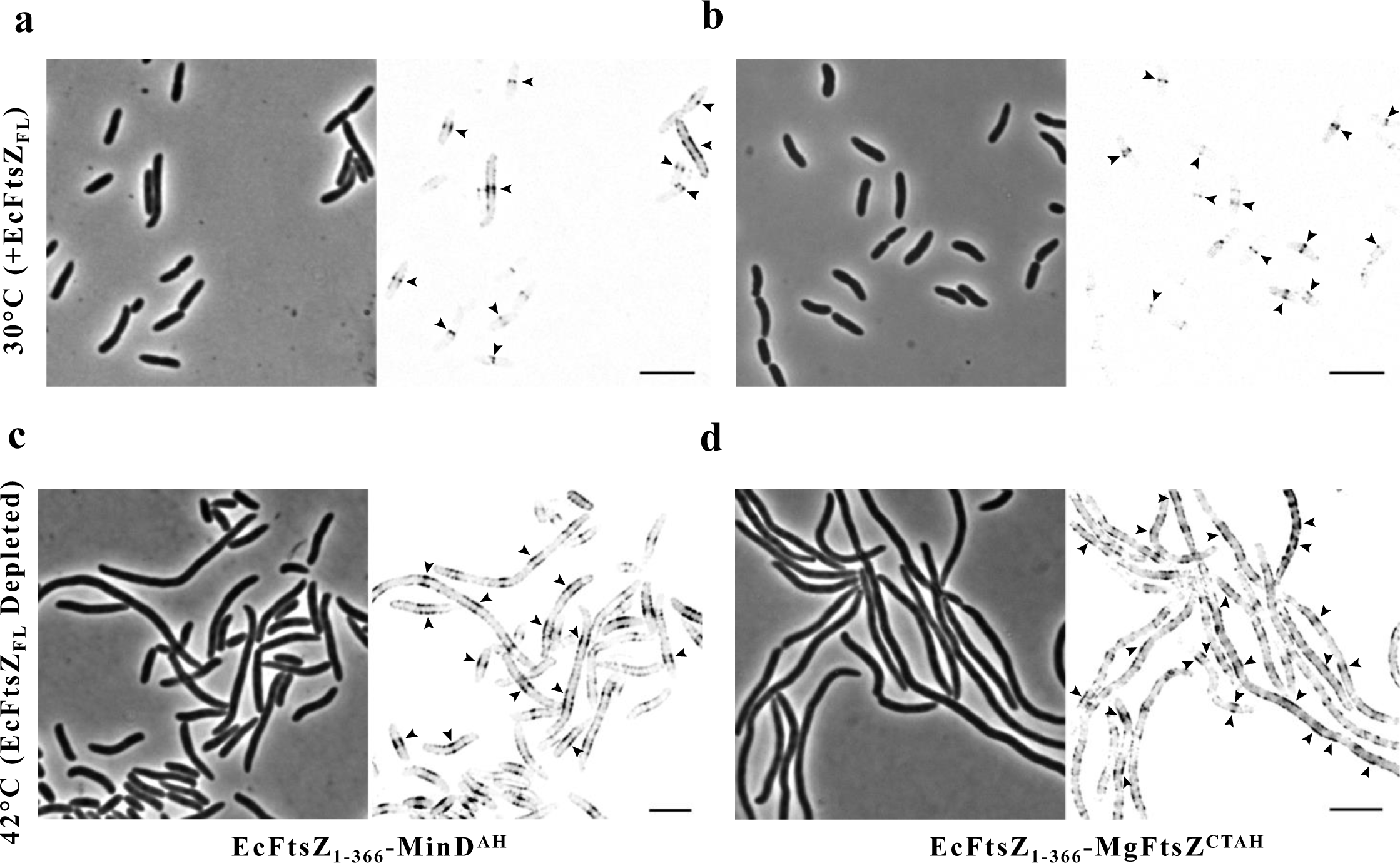
C-terminal amphipathic helix of MgFtsZ can bind to membrane *in vivo* in *E. coli* cells. The predicted C-terminal amphipathic helix of MgFtsZ can localise EcFtsZ_1-366_ (lacking CCTP) to membranes in *E. coli*. Localization of (i) EcFtsZ_1-366_-mNG-MinD^AH^ and (ii) EcFtsZ_1-366_-MgFtsZ^CTAH^ to Z-rings formed by endogenous native full-length FtsZ at 30 °C. Localization of (iii) EcFtsZ_1-366_-mNG-MinD^AH^ and (iv) EcFtsZ_1-366_-MgFtsZ^CTAH^ to inner membrane of filamentous cells due to depletion of native full-length FtsZ at 42 °C. Arrowheads mark the regions of membrane localisation. Scale bars represent 10 µm.

### Sequence analysis identifies a cholesterol recognition motif in *M. genitalium* and *M. pneumoniae* FtsZ

A careful analysis of the *M. genitalium* and *M. pneumoniae* FtsZ sequences identified a possible Cholesterol Recognition Amino acid Consensus (CRAC) motif located within the C-terminal helix (Fig. 5a). The CRAC motif is defined as (**L/V**)-X_1-5_-**Y**-X_1-5_-(**R/K**) where “X” represents any residue and central tyrosine has been shown to be essential for cholesterol binding (Jamin *et al.*, 2005). We found that *M. genitalium* contains CRAC sequence as **L^357^**-X-X-X-**Y^361^**-X-**K^363^** whereas *M. pneumoniae* has **L^367^**-X-X-X-**Y^371^**-X-**K^373^** in their respective C-terminal ends of FtsZs which may give specificity for binding to cholesterol and thereby to the membrane where 35-50% of total membrane lipid contents is cholesterol (47). The presence of CRAC motif was only found in *M. genitalium* and *M. pneumoniae* species with C-terminal amphipathic helix of the FtsZ sequences.

**Fig. 5.**
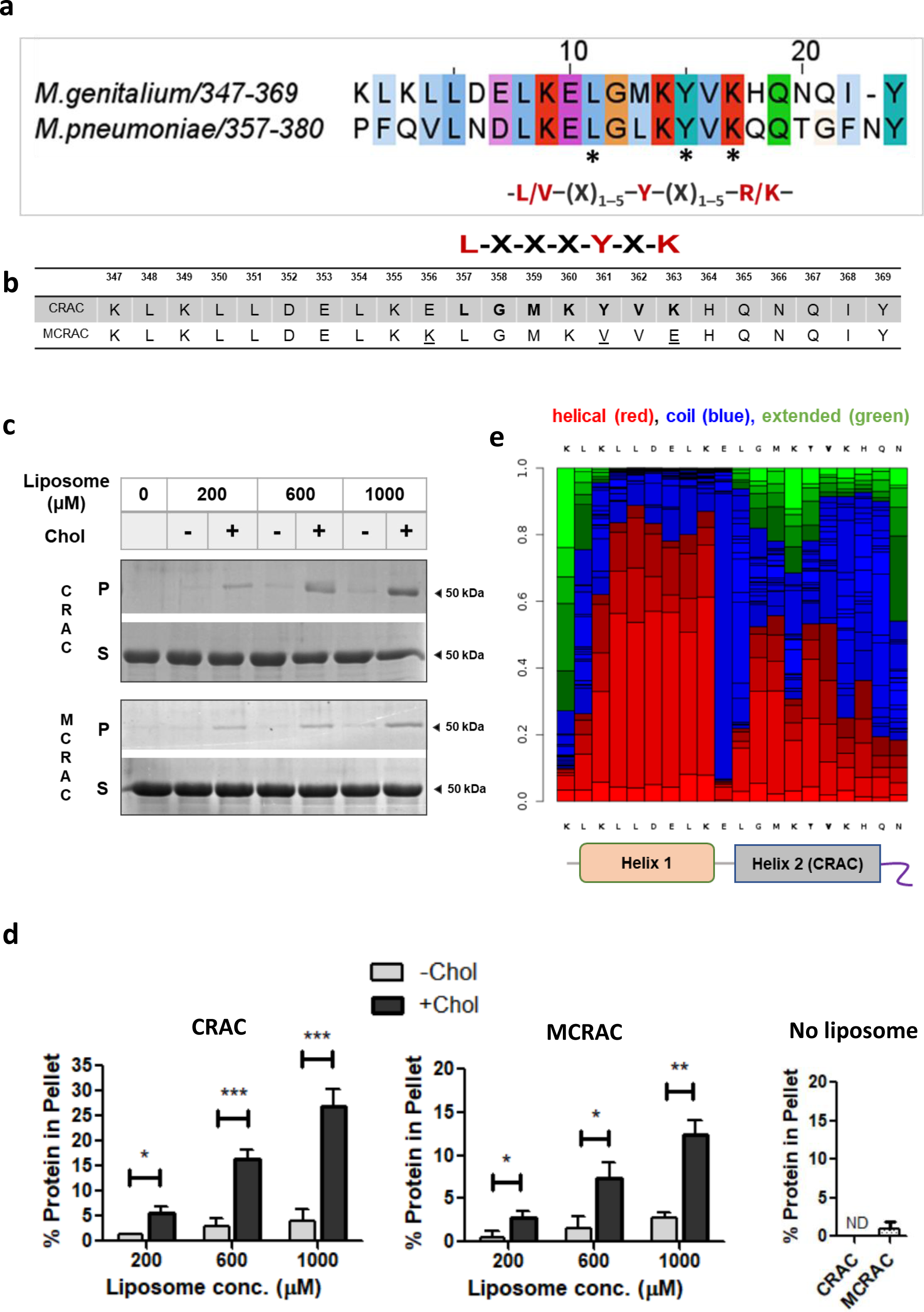
Cholesterol Recognition Amino acid Consensus (CRAC) motif in *M. genitalium* FtsZ provides it a preference for cholesterol rich liposome binding. (a) Sequence alignment of C-terminal end region of *M. genitalium* and *M. pneumoniae* FtsZ shows the presence of CRAC motif specific conserved residues (marked with ‘**\***’). The consensus motif is mentioned below the alignment. (b) Sequence of SmFtsZ^MgCTAH^ (denoted as CRAC) and SmFtsZ^MgCTAH(M-CRAC)^ (denoted as MCRAC) C-terminal motifs. The consensus amino acid sequence specific to CRAC motif are highlighted in bold. The mutated residues in MCRAC are underlined. (c) Representative 12% SDS-PAGE gels showing cholesterol mediated induction of membrane binding in CRAC (i.e SmFtsZ^MgCTAH^) and MCRAC (i.e. SmFtsZ^MgCTAH(M-CRAC)^) in liposome pelleting assay. Liposome containing cholesterol (+Chol) has higher amount of protein in pellet fraction compared to the liposome without cholesterol (-Chol). (d) Quantification of protein amount in pellet fraction in liposome pelleting assay. CTAH and MCRAC both show more binding with liposomes containing cholesterol with a significant difference in 1000 µM liposome conditions. Error bars indicate standard deviation from three independent experiments. p values are obtained from an unpaired two tailed t-test using the function ‘T.TEST’ in Microsoft Excel. Significance level is denoted by *, p < 0.05; **, p < 0.01; ***, p < 0.005); ND, not detected. (e) Local secondary structure of the C-terminal end region of *M. genitalium* FtsZ was predicted using PEP-FOLD server (https://bioserv.rpbs.univ-paris-diderot.fr/services/PEP-FOLD/). Two helical regions have been identified of which the Helix 2 contains the hypothetical CRAC motif sequence that could get inserted inside membrane to interact with cholesterol. The schematic shown below the plot represents the same.

To indeed confirm that the proposed CRAC motif confers a preference for cholesterol, we performed pelleting assay of SmFtsZ^MgCTAH^ construct with liposomes containing cholesterol (in molar ratio of 30% DOPG, 30% DOPC, 40% Cholesterol). We observed an increased membrane binding in cholesterol containing liposomes compared to liposomes made with 30% DOPG and 70% DOPC (Fig. 5c, 5d) which might indicate its preference for cholesterol. We then disrupted the proposed CRAC motif (MCRAC) by mutating three residues E356K, Y361V and K363E to see if it abolishes cholesterol specific binding (Fig. 5b). However, interestingly, SmFtsZ^MgCTAH(M-CRAC)^ too showed increased binding similar to SmFtsZ^MgCTAH^ in the liposome pelleting experiments (Fig. 5c, 5d). Based on the local secondary structure prediction of the C-terminal 23 amino acid peptide stretch, we hypothesise two helical regions to be present separated by a short break in the helix with residues having low helix forming probability (Fig. 5e) while the CRAC motif containing helix binds to cholesterol by getting inserted inside membrane. The mutant construct SmFtsZ^MgCTAH(M-CRAC)^ might result in increased binding because of the variation in membrane binding properties and it was challenging to delineate the distinction between the two functions (cholesterol recognition and membrane binding).

## Discussion

Here, we provide evidence that a phylogenetically separate FtsZ clade among mycoplasmas possesses an inherent amphipathic helix that might facilitate FtsA-independent membrane attachment. We experimentally show the putative C-terminal amphipathic helix of *M. genitalium* FtsZ binds liposomes. Also, when we expressed the chimeric protein in *E. coli* cell in a FtsZ depleted condition, it showed membrane localization which further confirms membrane binding ability of MgFtsZ^CTAH^ motif. In minimal genome bacterial species like mycoplasma, the possibility of direct membrane binding in cell division protein FtsZ, bypassing the requirement of additional adaptor proteins, is of relevance. FtsZ with a membrane binding helix provides an ideal minimal system for mechanistic insights into the process of cell constriction and division. Additionally, the presence of a CRAC motif makes this core division protein an interesting model for studying division in mycoplasmas.

In previous studies, based on protein sequence analysis, the presence of amphipathic helix was proposed in the N-terminal tail region of archaeal FtsZ (48). Another report on a novel species of *Atribacteria* predicted a similar amphipathic helical region at the N-terminal extension of FtsZ, which might bind to membrane (49). However, the membrane tethering role of the proposed amphipathic helices from FtsZ has not been experimentally demonstrated till date. Recently eukaryotic chloroplast FtsZ1 from *Arabidopsis thaliana* has been shown to interact with membranes via its C-terminal amphipathic β-strand motif (50). It is interesting that eukaryotic FtsZs have also evolved a mechanism of an inherent amphipathic helix directly binding to the membrane, making the anchoring role of FtsA redundant.

In the case of mycoplasma FtsZs, we found both N- and C-terminal putative membrane attachment sites. The functional significance of such oppositely oriented membrane binding sites is currently not clear from the perspective of FtsZ filament structure and mycoplasma cell division. Earlier, a chimeric construct of FtsZ fused to a membrane tethering sequence (from MinD protein) was shown to constrict tubular liposomes (51). It has been also shown that changing the position of membrane targeting sequence in FtsZ from the conventional C-terminal end to N-terminal end results in convex bulge rather than concave depression (52). Interestingly in both the cases bending force could be generated which lead to the tubulation of the liposomes *in vitro*. However, experiments with membrane targeting sequence attached with FtsZ protein did not show full constriction. Septation was only sometimes observed upon addition of FtsA indicating the requirement of FtsA and its role beyond Z-ring anchoring (53). Loose and Mitchison’s *in vitro* experiments with *E. coli* FtsZ and FtsA on supported lipid bilayer also suggested a dual antagonistic role of FtsA in anchoring and assembling FtsZ filaments as well as its rapid disassembly from the membrane (34). Similarly, variations in the stoichiometry of FtsA and FtsZ revealed that excess FtsA can effectively destabilize FtsZ polymers *in-vitro* (33). FtsA has also been suggested to be involved in the septation process and membrane remodelling by introducing local curvature in the inner membrane (29). Thus, FtsA could play multiple roles as membrane anchor as well as in remodelling and regulating FtsZ filament dynamics. In contrast to the above experiments where FtsA was used to anchor FtsZ to the membrane, the *M. genitalium* FtsZA proteins, as revealed from our experiment and sequence analysis, are both potentially capable of membrane binding and possibly work in a novel way to drive cell division. We believe *M. genitalium* and *M. pneumoniae* FtsA could potentially serve as a model to study its fundamental role beyond just FtsZ tethering.

*M. genitalium* and *M. pneumoniae* among mycoplasmas were additionally found to contain a CRAC motif within the C-terminal amphipathic helix region. Our in vitro results also showed increased membrane binding to cholesterol containing liposomes indicating the possibility of cholesterol recognition through CRAC. CRAC motif has been known to exist in numerous proteins spanning across eukaryotic and prokaryotic organisms (54–56). In bacteria, CRAC motif has been so far identified solely in pathogenic bacterial toxins that specifically bind to the cholesterol present in the membrane of eukaryotic host cells (57, 58). CRAC sequence containing peptides within bacterial leukotoxin has been shown previously to have higher affinity towards cholesterol and mutation in the CRAC motif abrogates binding (59). However, our results of SmFtsZ^MgCTAH(M-CRAC)^, the mutated construct for CRAC motif, showed unaffected membrane binding property similar to SmFtsZ^MgCTAH^ in the presence of cholesterol. SmFtsZ^MgCTAH^ bound to liposomes composed of 30% DOPG and 70% DOPC, although a marked increase in binding was observed in liposomes with 30% DOPG, 30% DOPC, and 40% cholesterol or mycoplasma polar lipid composition which consists of 25% DOPG, 20% DOPC, 30% cardiolipin, and 25% sphingomyelin. Moreover, our *in vivo* experiments show that SmFtsZ^MgCTAH^ bind *E. coli* membrane as well although mycoplasma and *E. coli* have very different membrane composition. We think it is possible that the CRAC mutated SmFtsZ^MgCTAH(M-CRAC)^ construct still shows increased membrane affinity in presence of cholesterol because membrane binding can be determined by multiple other factors such as overall lipid composition, hydrophobicity of AH, charge pattern, and lipid-amino acid interactions. For example, earlier studies on a eukaryotic sterol synthesis enzyme squalene monooxygenase showed similar results where cholesterol induced regulation remined unaffected even after disrupting the central critical tyrosine and phenylalanine residue of the CRAC and CARC motif respectively present within the membrane binding loop (60, 61). Thus, from our preliminary data we believe the unique CRAC motif present in *M. genitalium* FtsZ protein is functional and can interact with cholesterol. Additional data in future will shed more light into its cholesterol binding properties. Notably, the presence of a CRAC motif within FtsZ might be of significance as this could imply its membrane localization and subsequent role in cell division can be cholesterol dependent especially when 35-50% total mycoplasma membrane lipids is composed of cholesterol. We also checked if similar cholesterol recognition sequences are present in the other Mycoplasma FtsZs (containing N-terminal AH) and FtsAs (containing C-terminal AH) and did not find any. Thus, the membrane attachment of other AH containing FtsZ and FtsAs appear to follow separate mechanisms. Future *in vivo* studies will possibly give us more information about how membrane cholesterol composition affects cellular functions of mycoplasma divisome machinery.

Overall, our analysis highlights different possible modes of FtsZ-anchoring to the mycoplasma membrane, which are shown schematically in Fig. 6a-e. Group 1A FtsZ binds to FtsA in a similar mechanism as found in cell walled bacteria with its canonical CCTP (Fig. 6a). In some species where FtsA lacks the amphipathic helix, FtsZ ring probably needs SepF for membrane anchoring. One similar example can be found in mycobacteria where SepF acts as the membrane anchor for FtsZ and become an essential protein in this organism which lacks homologs of membrane tethers like FtsA or ZipA (62). Importantly, SepF mediated membrane binding possibly occurs through hydrophobic loops rather than amphipathic helix. Next, group 1B and 1C have deviant type of CCTPs for binding FtsA. However, because group 1B FtsAs do not posses a canonical amphipathic helix, its binding towards membrane is unclear (Fig. 6b, c). It is possible that smaller motifs of one or two amino acids can contribute to membrane binding, as has been observed for DivIVA (63) and bactofilins (64). Finally, group 1D and 1E do not have a consensus CCTP and experimental proof of whether the non-canonical CCTP binds with FtsA is not available (Fig 6d, e). Group 1E FtsZs with N- or C terminal amphipathic helix raises the possibility of direct membrane attachment making FtsA’s membrane tethering function dispensable. Based on our results here, we hypothesise that the C-terminal amphipathic helix in *M. genitalium* and *M. pneumoniae* FtsZ will help it to directly attach to the membrane. Once binding happens, the CRAC motif gets inserted into the lipid bilayer and binds to interact with cholesterol (Fig. 6e right). However, additional studies are required to test and verify this hypothesis. We believe our findings in mycoplasmas to be an interesting observation to set the stage for designing new set of experiments and looking into a novel membrane tethering role of FtsZ as well as to study individual and cooperative roles of FtsZ and FtsA in ring constriction in the absence of cell wall synthesis machinery.

**Fig. 6.**
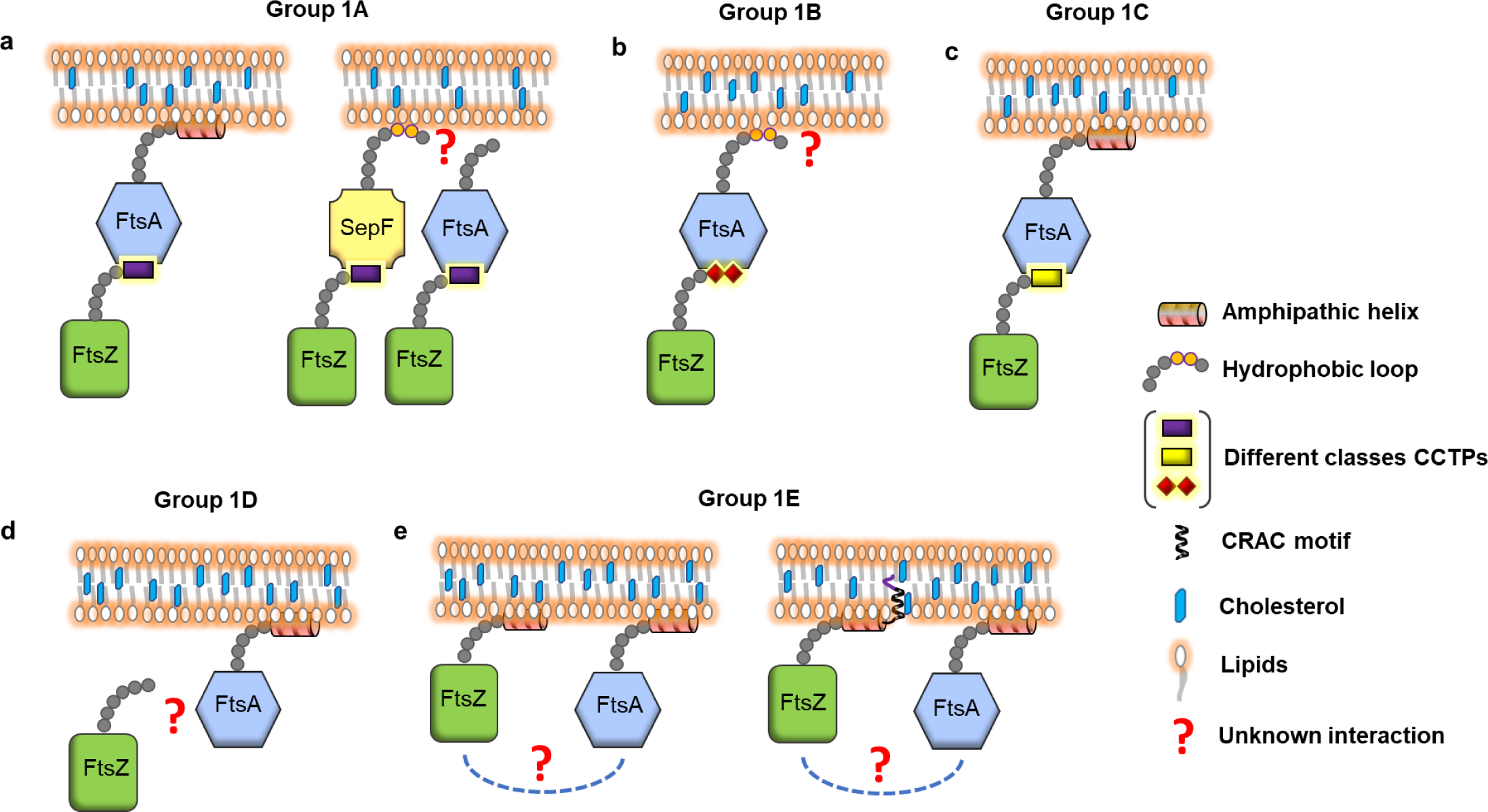
Modes of FtsZA interactions across different mycoplasma groups. The possible modes of FtsZ-FtsA interaction and their membrane-binding potentials are shown schematically for five mycoplasma groups.

## Materials and methods

### Sequence analysis of FtsZ and FtsA

We analysed a total 113 Mycoplasma species with complete sequence information of the FtsZ and FtsA from the NCBI genome database. The division and cell wall (*dcw*) cluster locus was first identified by manually checking genomes of every representative strain (detailed information of the strains is listed in supplementary data sheet) through a keyword-based search with ‘FtsZ’, ‘FtsA’, ‘rsmH’ (gene annotation for MraW protein) or ‘MraZ’. We had prior information about these four mycoplasma genes present in the dcw operon from the existing literature. We found that the upstream coding sequence to the *ftsZ* gene was annotated as a hypothetical protein-coding gene except for *Mycoplasma mycoides,* where it is annotated as *ftsA*. Indeed, in some earlier published literature, the hypothetical gene has been reported as ‘*ftsA*’ (65). Hence, we downloaded the protein sequences of the *ftsZ* gene along with the hypothetical protein-coding gene upstream of *ftsZ* for our analysis. We verified all the FtsA sequences by multiple sequence alignment with known and already characterized FtsA sequences from cell-walled model bacteria, followed by manual assignment of secondary structures present in FtsA. Multiple sequence alignment of FtsZ and FtsA sequences were performed in Jalview (https://www.jalview.org/) using Muscle (Edgar, 2004) with the default parameter. FtsA domain assignment was done in PROMALS3D (http://prodata.swmed.edu/promals3d/promals3d.php). The sequence logo was created using WebLogo (https://weblogo.berkeley.edu/logo.cgi). For secondary structure prediction of protein sequences, we used the PSIPRED server (http://bioinf.cs.ucl.ac.uk/psipred/). HeliQuest (https://heliquest.ipmc.cnrs.fr/cgi-bin/ComputParams.py) was used for helical wheel representation and calculation of hydrophobic moment and hydrophobicity parameters. Phylogenetic trees were constructed using maximum likelihood method in MEGA 11 (66) software with default parameters and visualized in iTOL (https://itol.embl.de/) (67).

### Cloning and protein purification

*SmftsZ* gene was PCR amplified from the chromosomal DNA of *Spiroplasma melliferum* KC3 followed by restriction digestion and ligation at the Nde1 and BamH1 sites into a (His)_6_ tag containing pHis17 vector as reported earlier (68). Restriction free cloning (69) was used for cloning of untagged SmFtsZ construct using specific primers. A detailed list of plasmid constructs and oligonucleotide primers used in our study is given in Table S3. SmFtsZ^MgCTAH^ was created by replacing 45 base pairs from the 3’ end of the untagged SmFtsZ sequence with respective 69 base pairs of *M. genitalium ftsZ* using PCR. The amplified product was then inserted into (His)_6_-SUMO-pET28a vector at the Nde1 and BamH1 sites by restriction digestion and ligation. The construction of pJSB2-EcFtsZ_1-366_-mNG-MinD^AH^ (pCCD907) will be described in detail elsewhere, but is identical to the EcFtsZ_1-366_-mVenus-MinD^AH^ described by Erickson and colleagues (52), except that mNeonGreen replaces mVenus. The EcMinD residues corresponding to amino acids 254 – 270 in pJSB2-EcFtsZ_1-366_-mNG-MinD^AH^ were replaced with the predicted C-terminal amphipathic helix (23 amino acids corresponding to 347 – 366) from MgFtsZ by restriction free cloning to give pJSB2-EcFtsZ_1-366_-mNG-MgFtsZ^CTAH^ (pCCD1071). pJSB2 is pBAD derivative carrying the tightly regulated arabinose inducible promoter (70).

Untagged SmFtsZ was expressed in *E. coli* BL21-AI cells by 0.2% arabinose induction at OD600= 0.6-0.8 and was grown at 30°C for 5 hours. Cells were harvested by centrifugation at 4,000xg at 4°C for 20[mins and cell pellet was flash-frozen in liquid nitrogen and stored at -80°C until processed. On the day of purification, cell pellet was taken out from -80°C and thawed in ice for 10 mins and then resuspended in lysis buffer (50 mM Tris-HCl pH 8.0, 50[mM NaCl, 10% glycerol) followed by sonication to lyse the cells. The total lysate was spun at 30,000xg for 1 hour to remove the cell debris and the clear supernatant was loaded into HiTrap Q HP 5 ml column (GE) equilibrated in Buffer A50 (50 mM Tris-HCl pH 8.0, 50[mM NaCl). Protein was eluted in 5 mL fractions using a linear KCl gradient from 50 mM to 1000 mM over 20 column volumes. Peak fractions containing proteins were confirmed from SDS PAGE, collected and pooled together for dialysis against A50 buffer and loaded into MonoQ column to increase the purity level. Finally, the pure fractions were confirmed from SDS PAGE gel and pooed for dialysis in 50 mM HEPES KOH storage buffer (pH 7.4, 50 mM KCl). After dialyzing for 2 hours, the protein was concentrated using centricon (with 10 kDa membrane cut off), flash-frozen in aliquots and stored at -80°C.

SmFtsZ^MgCTAH^ (in pET28a SUMO vector) was expressed in *E. coli* Rosetta (DE3) pLysS at OD600= 0.5-0.6. by inducing with 0.6 mM IPTG and grown at 18°C for 16 hours. Cells were harvested by centrifugation at 4,000xg for 20[mins at 4°C and cell pellet was flash-frozen in liquid nitrogen and stored at -80°C until purification. Cells were resuspended in lysis buffer (50 mM Tris-HCl pH 8.0, 200 mM KCl, 10% glycerol) and sonicated to lyse the cells. The lysate was centrifuged at 30,000×g at 4°C for 1 hour and the supernatant was loaded to a 5 mL His-Trap column (GE), equilibrated with Buffer A (50 mM Tris-HCl pH 8.0, 200 mM KCl). Protein was eluted using a step gradient of 2%, 5%, 10%, 20%, 50% and 100% Buffer B (50 mM Tris-HCl pH 8.0, 200 mM KCl, 500 mM Imidazole). To cleave the N-terminal (His)_6_-SUMO tag, peak protein fractions were pooled and Ulp1 protease was added in a 1:100 (protease:protein) molar ratio and dialysis was set up against the cleavage buffer (50 mM Tris-HCl pH 8.0, 200 mM KCl, 0.5 mM DTT) for 10 hours at 4°C. (His)_6_-SUMO tag and uncleaved protein was removed by passing the dialysed sample mixture through a His-Trap column and flowthrough containing the SmFtsZ^MgCTAH^ protein was collected. Finally, the protein was dialysed in 50 mM HEPES KOH storage buffer (pH 7.4, 50[mM KCl) for 2 hours, concentrated using centricon (with 10 kDa membrane cut off), flash-frozen in aliquots and stored at -80°C. SmFtsZ^MgCTAH(M-CRAC)^ protein purification was done using the same protocol as SmFtsZ^MgCTAH^. All three purified proteins were run on a 12% SDS PAGE for checking the purity level and quality (Fig. S8).

### Growth conditions and microscopy

*E. coli* strains were grown in LB medium (Lennox or Miller) at 30 °C containing antibiotics as required. Carbenicillin, kanamycin and chloramphenicol were used at 100 µg/ml, 25 µg/ml and 34 µg/ml, respectively. Ampicillin was used at 200 µg/ml. *E. coli* strain JKD7_1 (W3110 *recA56 ftsZ::kan*) carrying pKD3 (Amp^R^, *repA^ts^*) (CCD56; (70) and the plasmid pJSB2 (Cm^R^) (pCCD434; Stricker and Erickson, 2003), pJSB2-EcFtsZ_1-366_-mNG-MinD^AH^ (pCCD907) or pJSB2-EcFtsZ_1-366_-mNG-MgFtsZ^CTAH^ (pCCD1071) was grown in LB Lennox at 30 °C (permissive temperatures) or at 42 °C (non-permissive temperature) to allow for the loss of pKD3 plasmid. Expression of EcFtsZ_1-366_-mNG-MinD^AH^ and EcFtsZ_1-366_-mNG-MgFtsZ^CTAH^ from pJSB2 derived plasmids was achieved by the addition of 0.2 % L-arabinose. Repression was achieved by addition of 2% glucose.

For microscopy, JKD7-1/pKD3 strain carrying the various pJSB2 derived plasmids were grown in repression media (2 % glucose) at 30 °C and diluted 1:100 into fresh LB Lennox medium containing 0.2 % L-arabinose and grown further at 42 °C for 2.5 hours. The cultures were grown to an OD_600_ between 0.4 - 0.6 and 1.5 mL culture was centrifuged at 1400 x *g* for 3 minutes. The cells pellets were resuspended in 100 – 150 µL of fresh LB medium, mounted on LB-agarose pads (1.6 %) and imaged using an epifluorescence microscope (DeltaVision^TM^Elite) with a 100X oil immersion phase objective (PLN100XOPH) of NA 1.25. and equipped with an in-line CCD camera (CoolSNAP^TM^ HQ2). Solid-state illumination (Spectra 7, Lumencor Light Engine^©^) was used as a light source with a 1 mm diameter liquid light guide. Excitation filters and emission filters of 488/28 nm and 525/48 nm respectively were used for imaging mNeonGreen tagged strains. Images were processed using Fiji v2.9.0/ 1.53t (71).

### Liposome preparation and sedimentation assay

Liposomes were prepared by using synthetic lipids DOPC (1,2-dioleoyl-sn-glycero-3-phosphocholine), DOPG (1,2-dioleoyl-snglycero-3-phospho-rac-1-glycerol), and DPG/cardiolipin (1,3-bis[1,2-dioleoyl-sn-glycero-3-phospho]-glycerol) and SM (sphingomyelin), all purchased as chloroform solution from Avanti Polar Lipids, Inc. Cholesterol powder was purchased from Sigma. To prepare 2 mM stock liposomes, the required volume of individual components (as per molar ratio) was first taken in a glass tube. The chloroform was evaporated under a stream of nitrogen and further kept for drying at room temperature for 15 mins. A thin layer of lipids was visible at the bottom wall of the tube once the chloroform was dried. The dried lipid mixture was hydrated with the required volume of reaction buffer containing 50 mM HEPES KOH, 50 mM KCl (pH 7.4). After vortexing well, the mixture became turbid and it was extruded 15-20 times through a 100 nm pore-sized polycarbonate membrane until the solution became translucent.

For liposome sedimentation assay, purified flash-frozen protein was taken out from -80°C, thawed in ice for 10 mins and spun at 21,000xg for 1 hour at 4°C to remove any precipitation and supernatant was used in the assay. We set up a 100 µL reaction with 2 µM of protein and a varying concentration of liposome from 0 µM (i.e., no liposome control) to 1000 µM. The reaction mixture was incubated at 30°C for 20 mins and spun at 21,000xg for 40 mins to pellet down the liposomes (for pelleting cholesterol containing liposomes, ultracentrifugation at 100,000xg was used for 30 mins). Immediately after the spin, we took out the supernatant, washed the pellet with 100 µL of buffer and resuspended it in the same volume. Next, the supernatant and the pellet samples were mixed with 2X Laemmli buffer and equal amounts of both were loaded and run into a 12% SDS-PAGE gel followed by staining with Coomassie Brilliant Blue to check the presence of protein on the gel. Experiments were performed in triplicates and representative gel images are shown. The percentage intensities of protein bands in pellet and supernatant fractions were calculated as intensity of pellet (or supernatant) divided by sum of the two intensities. The intensity of the bands were quantified using the ImageJ software (Rueden *et al.*, 2017) by measuring the area of the respective bands and percentage of protein in pellet fractions were calculated by dividing the pellet band intensity by the sum of the pellet and supernatant band intensity. All the plots were generated in GraphPad Prism software.

## Author Contributions

SD and PG conceived the project and designed the experiments. SD conducted all experiments and analysed the data except mentioned otherwise. JC contributed to cloning and protein purification of untagged SmFtsZ. SMP carried out and analysed the *in vivo* experiments related to imaging of JKD7-1 strains, while RS supervised, analysed the work and edited the manuscript. SD and PG wrote the manuscript. PG supervised the project and acquired funding. All authors read and approved the final submitted version of the manuscript.

## Acknowledgments

All authors sincerely thank IISER Pune for all the facilities to carry out this research. We thank Ajay Kumar Sharma for the construction of pJSB2-EcFtsZ1-366-mNG-MinD^AH^. We thank Harold Erickson (Duke University, USA) for strains and plasmids. We acknowledge fellowships from CSIR to SD, IISER Pune to JC and NISER, DAE to SMP. Research work in the lab of PG was supported by Department of Biotechnology (DBT) Membrane Structural Biology Program grant (BT/PR28833/BRB/10/1705/2018), SERB CRG (CRG/2018/003795) and IISER Pune. Work in RS lab was supported by SERB CRG (CRG/2021/000337) and intra-mural funding from Department of Atomic Energy (DAE).

