## Supplemental figures and tables for "An amphipathic helix facilitates direct membrane binding of Mycoplasma FtsZ"

**Table S1: Domain boundaries of Group 1E Mycoplasma FtsZ sequences with amphipathic helix**

| Organism | Conserved domains | | | |
|---|---|---|---|---|
| | N-term extension (AH) | Globular core | Linker | C-term end (AH)* |
| Candidatus Mycoplasma girerdii | 1-43 (3-12) | 44-352 | 353-441 | 442-452 |
| Mycoplasma alvi | 1-115 (7-25) | 116-423 | 424-532 | 533-542 |
| Mycoplasma amphoriforme | 1-137 (6-17) | 138-451 | 452-562 | 563-573 |
| Mycoplasma gallisepticum | 1-74 (11-37) | 75-378 | 379-444 | 445-455 |
| Mycoplasma imitans | 1-77 (11-32) | 78-382 | 383-448 | 449-459 |
| Mycoplasma iowae | 1-121 (5-14) | 122-428 | 429-545 | 546-554 |
| Mycoplasma penetrans | 1-56 (26-47) | 57-364 | 365-498 | 499-509 |
| Mycoplasma pirum | 1-94 (2-12) | 95-401 | 402-505 | 506-516 |
| Mycoplasma testudinis | 1-153 (4-12) | 154-465 | 466-574 | 575-585 |
| Mycoplasma tullyi | 1-79 (13-33) | 80-383 | 384-449 | 450-460 |
| Mycoplasma genitalium | 1-18 | 19-327 | 327-346 | 347-369 (347-366)* |
| Mycoplasma pneumoniae | 1-18 | 19-329 | 330-353 | 354-380 (354-375)* |

**Table S2: Details of amphipathic helical parameters of NTAH and CTAH sequences of Group 1E *Mycoplasma* FtsZ sequences**

| Organism | NTAH/CTAH sequence | Hydrophobicity | Hydrophobic moment |
|---|---|---|---|
| *Candidatus Mycoplasma girerdii* | INKILDEIIN | 0.530 | 0.857 |
| *Mycoplasma alvi* | MKDVLSSFDSDLSQDFLKT | 0.313 | 0.376 |
| *Mycoplasma amphoriforme* | RMNDALFHFDE | 0.287 | 0.417 |
| *Mycoplasma gallisepticum* | KSLEEMQEELIKLQKEMQAVLM KDRA | 0.173 | 0.4 |
| *Mycoplasma imitans* | KSLEAMQEQLIKLQKEMEAVLM | 0.359 | 0.465 |
| *Mycoplasma iowae* | RSNIDKILNS | 0.125 | 0.541 |
| *Mycoplasma penetrans* | IKKEINILQEINNSTLVRLLSN | 0.387 | 0.321 |
| *Mycoplasma pirum* | MKKYNQMQEVLS | 0.220 | 0.445 |
| *Mycoplasma testudinis* | LNRMNQVLF | 0.579 | 0.522 |
| *Mycoplasma tullyi* | LEAMQEELIKLQKEMQAELMK | 0.290 | 0.473 |
| *Mycoplasma genitalium* | KLKLLDELKELGMKYVKHQ | 0.254 | 0.381 |
| *Mycoplasma pneumoniae* | QVLNDLKELGLKYVKQQ | 0.268 | 0.432 |

**Table S3: Construct and plasmid details**

| Oligo name | 5'-3' sequence | Construct made | Vector backbone | Reference |
|---|---|---|---|---|
| SmFtsZ_vF | GTTTAACTTTAAGAAGGAGATATACATATGGACAATTTTGATAATTATG | Untagged SmFtsZ | pHis17-$Amp^R$ | (68) |
| SmFtsZ_stop_vR | GCTTTTAATGATGATGATGATGATGGGATCCTTACCAACTACGACGAACAAATG | Untagged SmFtsZ | pHis17-$Amp^R$ | This work |
| SmFtsZ-CTAH-RP1 | CTTAACATATTTCATGCCAAGTTCTTTCAGCTCATCTAAAAGTTTCAACTTATGTTCAACTACTTGATTATTA | $SmFtsZ^{MgCTAH}$ | pET28a-$Kan^R$ | This work |
| SmFtsZ-CTAH-RP2 | GCTTTTAATGATGATGATGATGATGGGATCCGTAGATTTGGTTTTGGTGCTTAACATATTTCATGCCAAGTTC | $SmFtsZ^{MgCTAH(M-CRAC)}$ | pET28a-$Kan^R$ | This work |
| MCRAC Rev1 | GGTTCTGATGTTCAACAACTTTCATGCCAAGTTTCTTCAGCTCATC | $SmFtsZ^{MgCTAH(M-CRAC)}$ | pET28a-$Kan^R$ | This work |
| MCRAC Rev2 | CTCGAATTCGGATCCTCAATAAATCTGGTTCTGATGTTCAACAAC | $SmFtsZ^{MgCTAH(M-CRAC)}$ | pET28a-$Kan^R$ | This work |
| mNG CTAH RP1 | ATTTCATGCCAAGTTCTTTCAGCTCATCTAAAAGTTTCAACTTACCACCACCCTTGTACAGTTCG | $EcFtsZ_{1-366}$-mNG-$MgFtsZ^{CTAH}$ | pJSB2-$Cm^R$ | This work |
| CTAH pJSB RP2 | CCGCCAAAACAGCCAAGCTTAATAAATCTGGTTCTGATGTTTCACATATTTCATGCCAAGTTCTTTC | $EcFtsZ_{1-366}$-mNG-$MgFtsZ^{CTAH}$ | pJSB2-$Cm^{Rs}$ | This work |
| mNG Forward | GTCAGCAAAGGTGAAGAAGACAATATGG | $EcFtsZ_{1-366}$-mNG-$MgFtsZ^{CTAH}$ | pJSB2-$Cm^R$ | This work |

**Fig. S1**

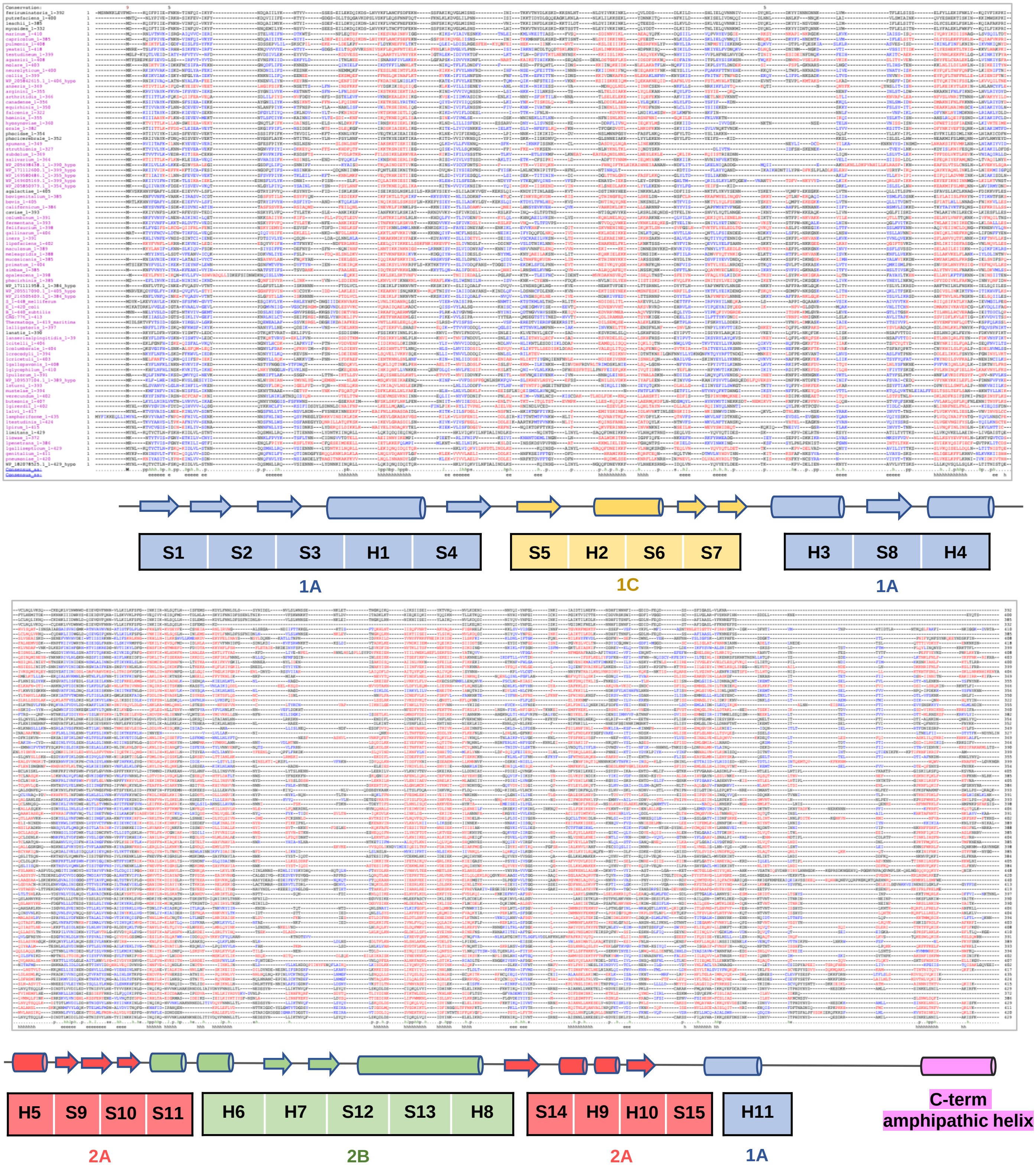


### Fig. S1 Domain assignment of hypothetical FtsA proteins in mycoplasma

Multiple sequence alignment of hypothetical protein sequences upstream of FtsZ using PROMALS 3D. The red and blue color code in each sequence represents helix and strand secondary sequences respectively. The secondary structures and domain assignment of FtsA are shown below the alignment showing the presence of canonical FtsA domain architecture.

Fig. S2

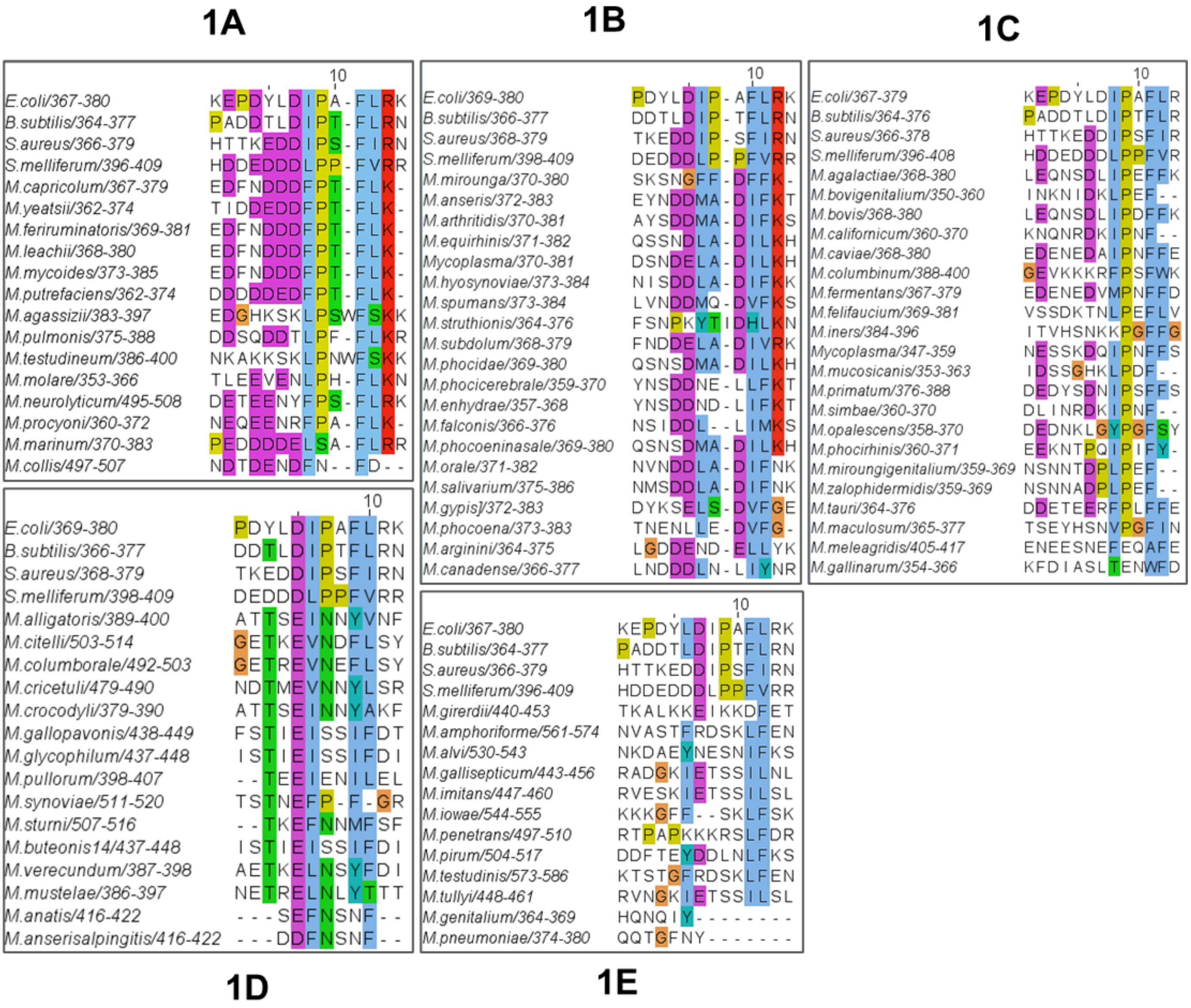


**Fig. S2 Sequence alignment of CCTP region of FtsZ proteins form group 1 species of mycoplasma.**

Various types of CCTPs are found across five mycoplasma groups. While group 1A-1C CCTPs show more similarity to the conserved canonical CCTP of cell-walled bacteria, groups 1D and 1E do not. The alignment of CCTP motifs shown here is used to generate the WebLogos in Fig. 2a.

**Fig. S3**

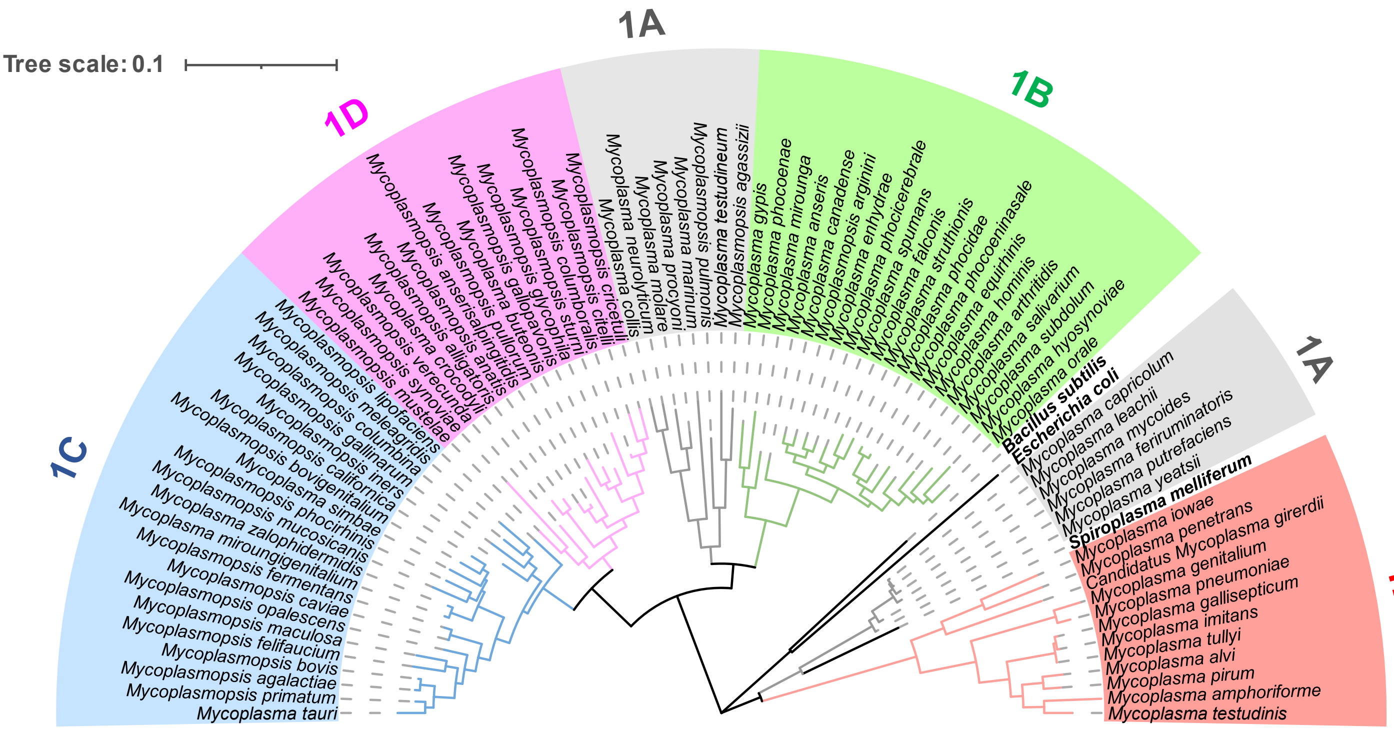


**Fig. S3 16S rRNA based cladogram of group 1 mycoplasma species.**

A cladogram based on the 16S rRNA sequences out of group 1 species is constructed using the maximum likelihood method with default parameters in MEGA11 software.

Fig. S4

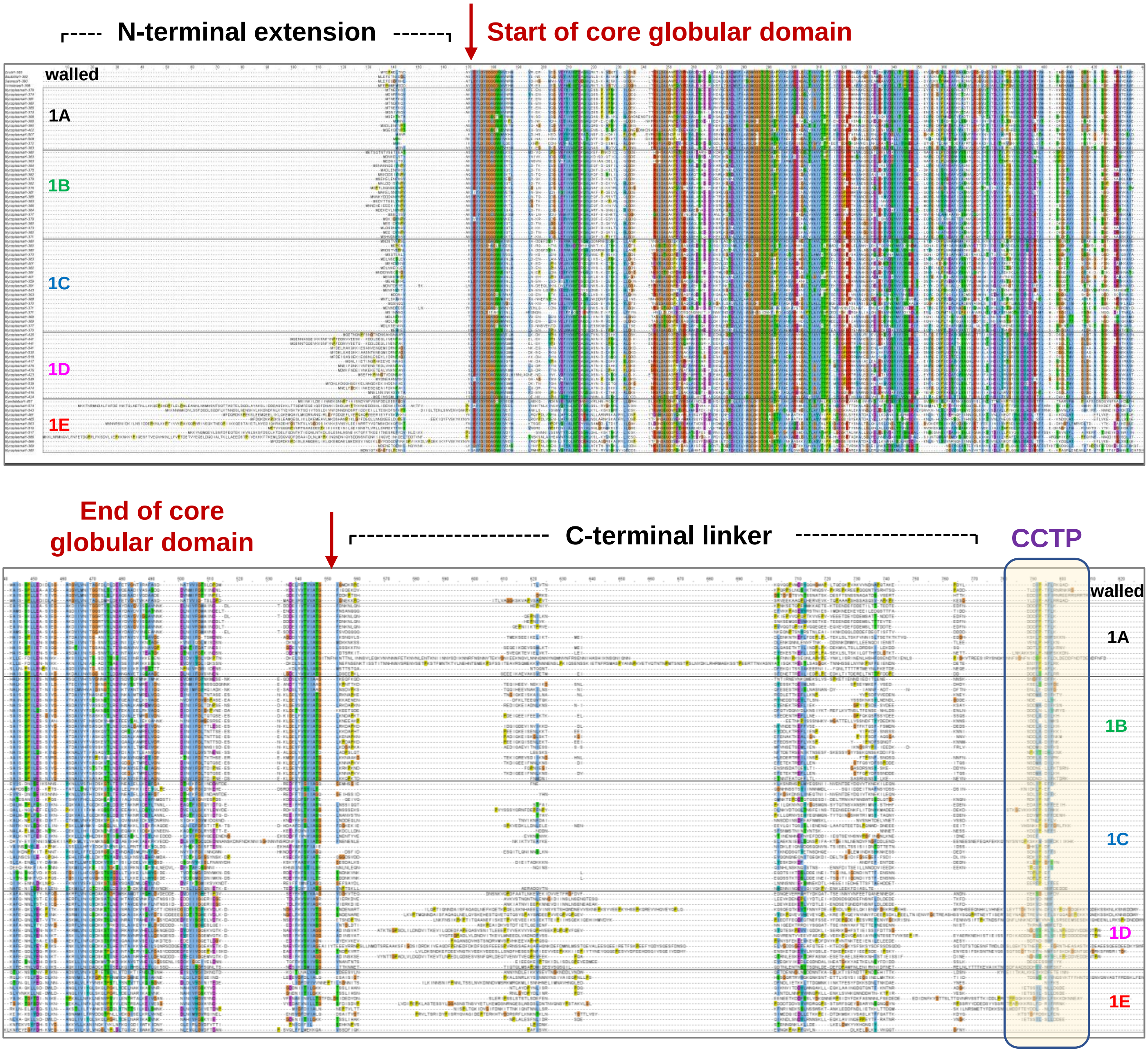


**Fig. S4 Sequence alignment of FtsZ proteins form group 1 species of mycoplasma.**

Multiple sequence alignment of FtsZ sequences shows the differences in the conservation at the C-terminal end compared to cell-walled bacterial FtsZs. Arrow indicates start and end of the globular core domain followed by a C-terminal disordered linker and CCTP. The C-terminal peptide region is highlighted within the yellow box. Multiple sequence alignment of FtsZ sequences and its homologs from cell-walled bacteria shows the presence of a longer N-terminal extension region in group 1D and 1E organisms.

**Fig. S5**

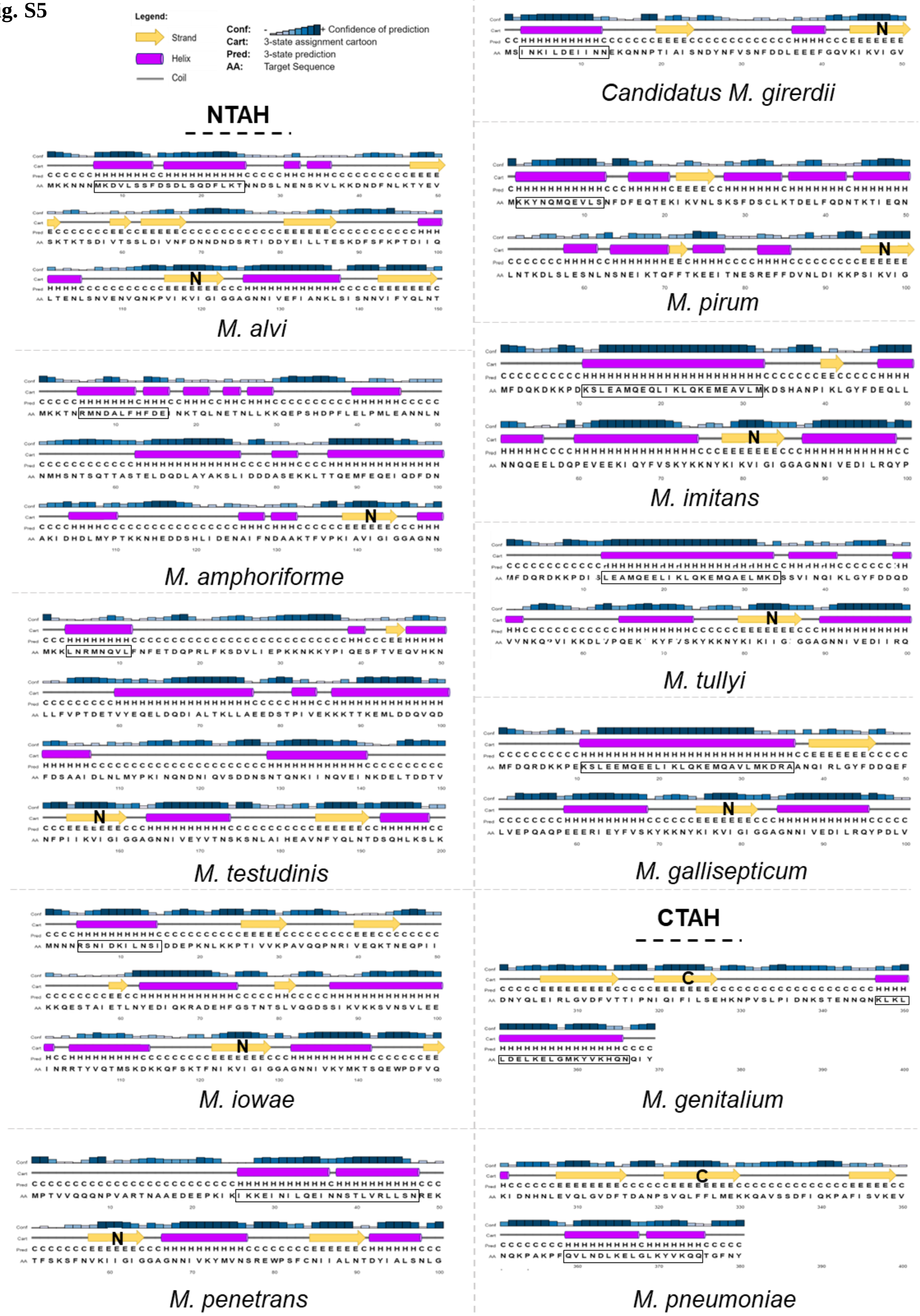
Legend:
Strand
Helix
Coil
Conf: Confidence of prediction
Cart: 3-state assignment cartoon
Pred: 3-state prediction
AA: Target Sequence
NTAH
M. alvi
M. amphoriforme
M. testudinis
M. iowae
M. penetrans
Candidatus M. girerdii
M. pirum
M. imitans
M. tullyi
M. gallisepticum
CTAH
M. genitalium
M. pneumoniae

**Fig. S5 Prediction of secondary structures of N- and C-terminal extensions of FtsZ in group 1E mycoplasma species.**

N-terminal amphipathic helix (NTAH) and C-terminal amphipathic helix (CTAH) containing FtsZ sequences are shown with secondary structures as predicted through PSIPRED. Helical regions with potential amphipathic nature are highlighted within the black rectangular box. The start and end of the globular domain of FtsZ are marked as ‘N’ and ‘C’ respectively.

**Fig. S6**

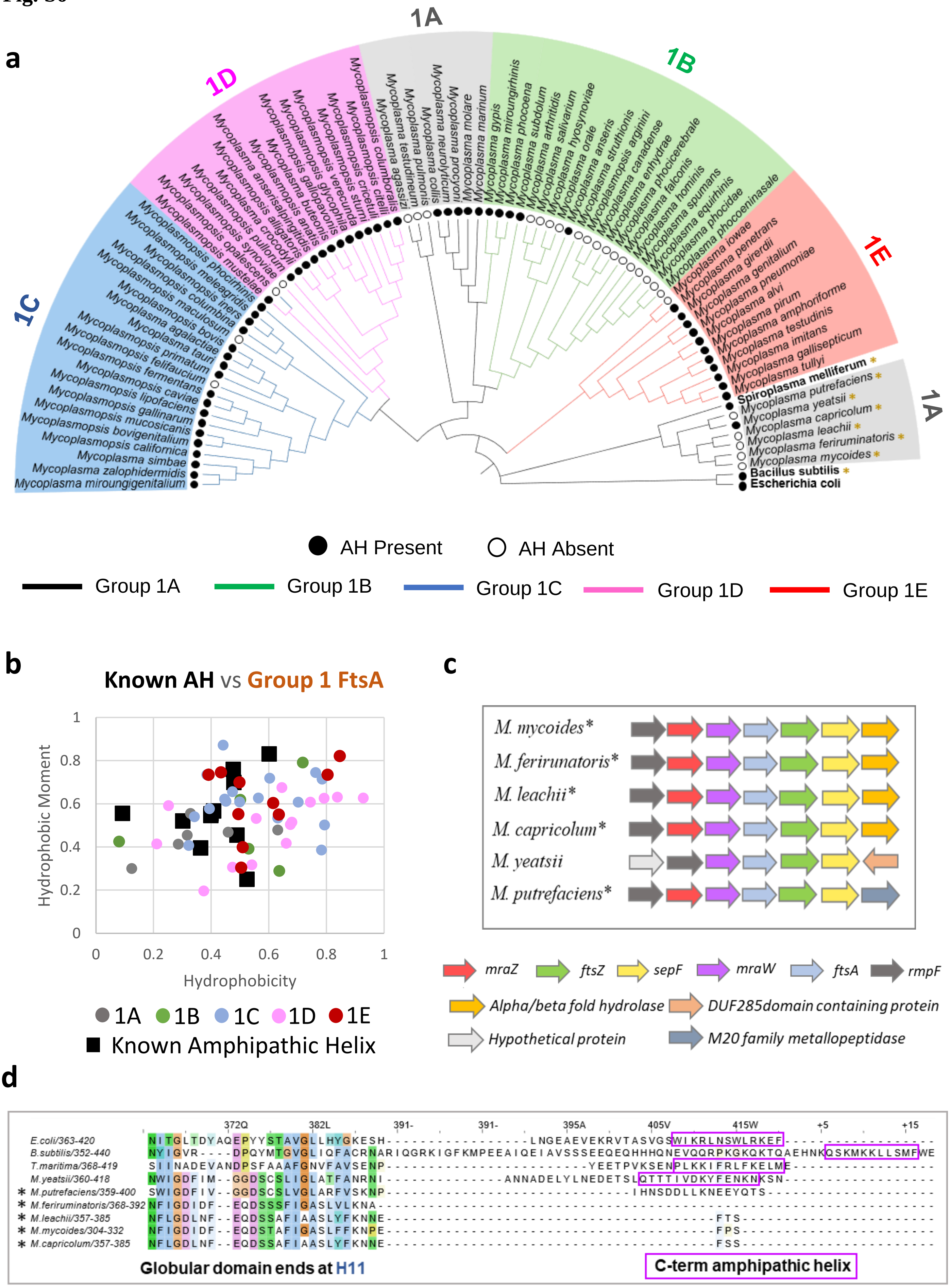
a
1A
1B
1C
1D
1E
Spiroplasma melliferum
Mycoplasma putrefaciens
Mycoplasma yeatsii
Mycoplasma capricolum
Mycoplasma leachii
Mycoplasma feriruminatoris
Mycoplasma mycoides
Bacillus subtilis
Escherichia coli
AH Present
AH Absent
Group 1A
Group 1B
Group 1C
Group 1D
Group 1E
b
Known AH vs Group 1 FtsA
Hydrophobic Moment
Hydrophobicity
1A
1B
1C
1D
1E
Known Amphipathic Helix
c
M. mycoides*
M. ferirunatoris*
M. leachii*
M. capricolum*
M. yeatsii
M. putrefaciens*
mraZ
ftsZ
sepF
mraW
ftsA
rmpF
Alpha/beta fold hydrolase
DUF285domain containing protein
Hypothetical protein
M20 family metallopeptidase
d
E.coli/363-420
B.subtilis/352-440
T.maritima/368-419
M.yeatsii/360-418
M.putrefaciens/359-400
M.feriruminatoris/368-392
M.leachii/357-385
M.mycoides/304-332
M.capricolum/357-385
Globular domain ends at H11
C-term amphipathic helix

**Fig. S6 Possibilities of non-FtsZ mediated modes of membrane attachment for the FtsZ ring across group 1 mycoplasma species.**

(a) A cladogram based on the FtsA sequences is constructed using the maximum likelihood method. The presence and absence of CTAH in FtsAs across mycoplasma species are depicted as filled and open circles respectively. The branches of the organisms are colored according to their respective groups. Mycoplasma species have SepF protein are marked with yellow colored star (*).

(b) Hydrophobic moment plot of all putative amphipathic helix mycoplasma FtsA from group 1. Known amphipathic helices from FtsA, MinD, and SepF are also plotted simultaneously for comparison.

(c) Schematic representation of dcw cluster genes in group 1A mycoplasmas containing SepF. Neighbouring genes (+1 and −1 positions) of dcw operon genes are also marked. The colour code and annotations are mentioned. The direction of the arrows indicates gene orientation 5′ → 3′. Organisms marked as '*' do not posses an amphipathic helix in their FtsAs.

(d) Sequence alignment FtsA showing the absence of a C-terminal amphipathic helix for organisms marked as black colored star (*****). Residues within pink colored box represent canonical C-terminal amphipathic helix in cell-walled bacteria.

**Fig. S7**

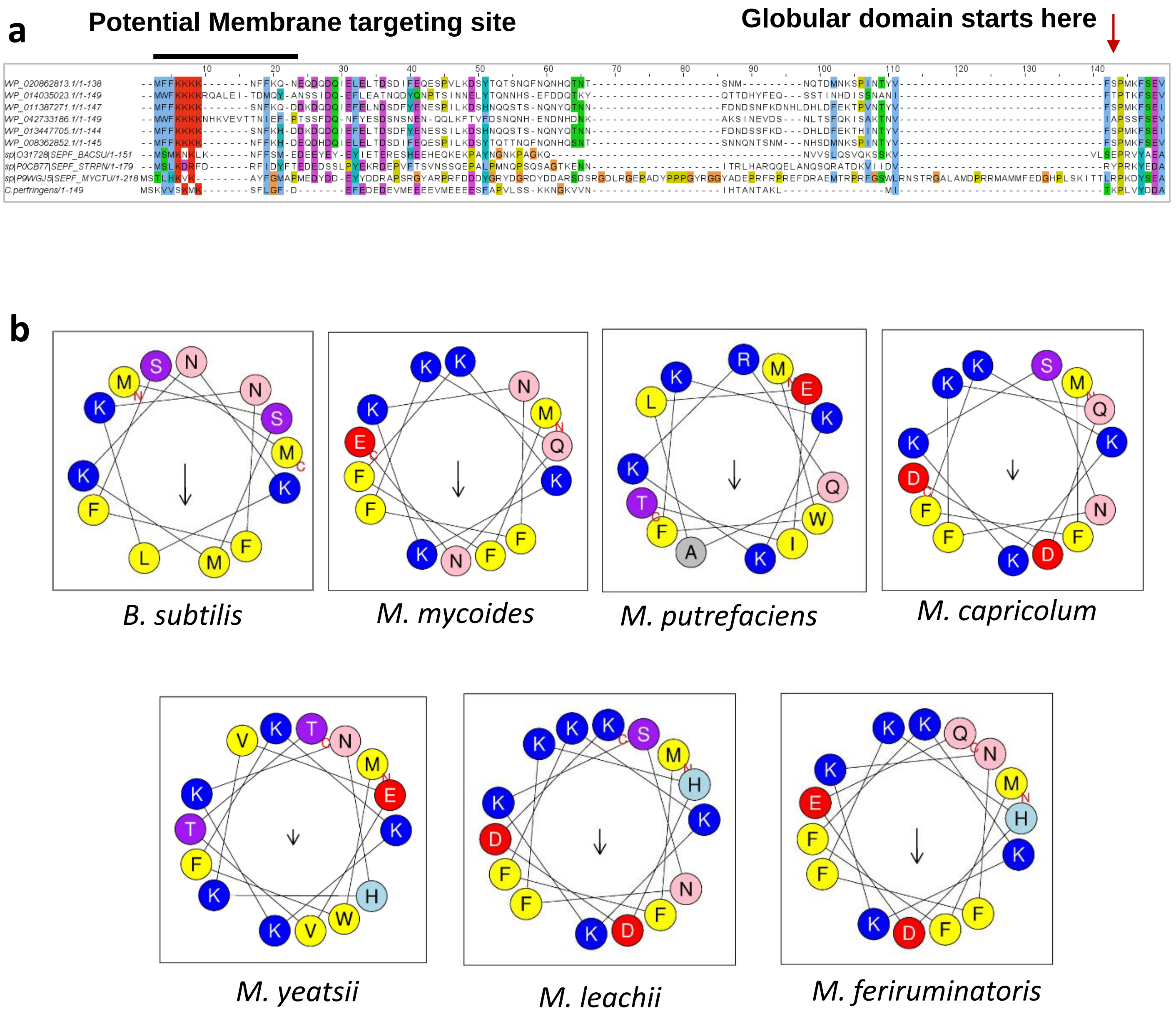


**Fig. S7 Prediction of amphipathic helix in mycoplasma SepF**

(a) Multiple sequence alignment of mycoplasma SepF proteins showing potential membrane binding region at the N-terminal end similar to what is present in *B. subtilis*, *S. pneumoniae*, *M. tuberculosis* and *C. perfringens* SepF.

(b) Helical wheel diagram of predicted membrane binding sites showing the presence of hydrophobic residues. Known membrane binding regions of SepF in *B. subtilis* has been included for comparison.

**Fig. S8**

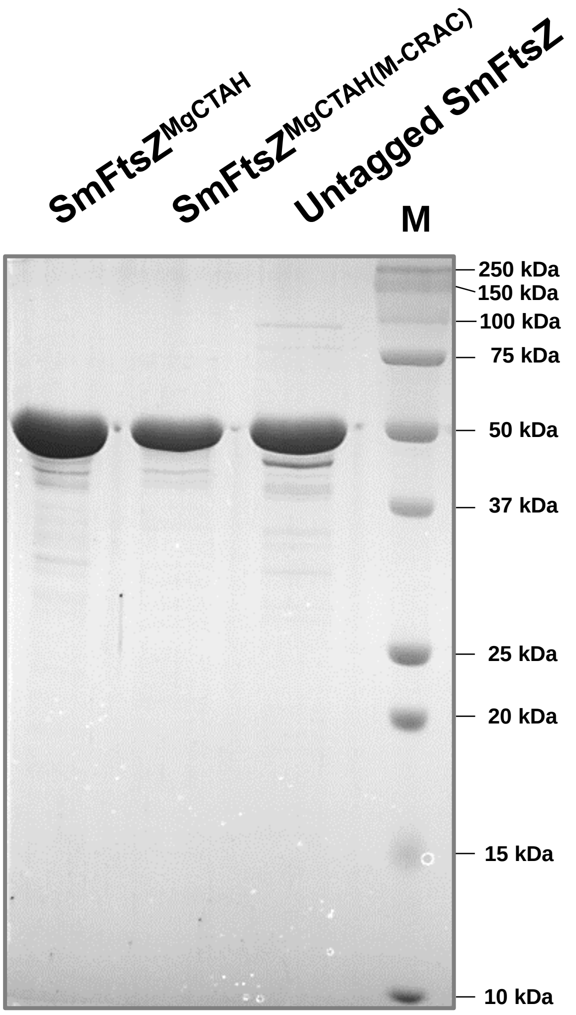


**Fig. S8 A representative gel showing the purity of purified proteins**

Purified proteins used in the study SmFtsZ$^{\mathrm{MgCTAH}}$, SmFtsZ$^{\mathrm{MgCTAH(M\text{-}CRAC)}}$ and Untagged SmFtsZ were run on a 12% SDS-PAGE gel. M lane corresponds to Protein ladder.

**Supplementary data sheet: Details of organism/strain information and accession numbers of proteins used for analysis**
